# Benchmarking clustering, alignment, and integration methods for spatial transcriptomics

**DOI:** 10.1101/2024.03.12.584114

**Authors:** Yunfei Hu, Yikang Li, Manfei Xie, Mingxing Rao, Wenjun Shen, Can Luo, Haoran Qin, Jihoon Baek, Xin Maizie Zhou

## Abstract

Spatial transcriptomics (ST) is advancing our understanding of complex tissues and organisms. However, building a robust clustering algorithm to define spatially coherent regions in a single tissue slice, and aligning or integrating multiple tissue slices originating from diverse sources for essential downstream analyses remain challenging. Numerous clustering, alignment, and integration methods have been specifically designed for ST data by leveraging its spatial information. The absence of benchmark studies complicates the selection of methods and future method development. Here we systematically benchmark a variety of state-of-the-art algorithms with a wide range of real and simulated datasets of varying sizes, technologies, species, and complexity. Different experimental metrics and analyses, like adjusted rand index (ARI), uniform manifold approximation and projection (UMAP) visualization, layer-wise and spot-to-spot alignment accuracy, spatial coherence score (SCS), and 3D reconstruction, are meticulously designed to assess method performance as well as data quality. We analyze the strengths and weaknesses of each method using diverse quantitative and qualitative metrics. This analysis leads to a comprehensive recommendation that covers multiple aspects for users. The code used for evaluation is available on GitHub. Additionally, we provide jupyter notebook tutorials and documentation to facilitate the reproduction of all benchmarking results and to support the study of new methods and new datasets (https://benchmarkst-reproducibility.readthedocs.io/en/latest/).

## Introduction

Spatial transcriptomics (ST) technology, emerging as a complementary approach to scRNA-seq, facilitates comprehensive gene expression profiling in tissue samples while preserving the spatial information of every cell or spot analyzed [1, 2]. ST techniques have significantly enhanced our understanding of cellular heterogeneity and tissue organization, offering insights into developmental processes, disease mechanisms, and potential therapeutic strategies [3–6]. Generally, ST technologies are commonly categorized into two groups: imaging-based and sequencing-based methods [7–13]. Continual progress in multiple ST technologies, including advancements in spatial resolution, capture capabilities, and computational methods, is continuously enhancing their potential applications and capabilities. An essential initial step in ST research is to cluster the spots and define spatially coherent regions in terms of expression data and location adjacency [14,15]. This process essentially entails classical unsupervised clustering of spots into groups according to the similarity of their gene expression profiles and spatial locations, subsequently assigning labels to each cluster. To date, existing clustering methods in ST can be broadly categorized into two main groups: statistical methods and graph-based deep learning methods [16].

Statistical models typically provide deterministic outputs; four representative methods are BayesSpace [17], BASS [18], SpatialPCA [19], and DR-SC [20]. BayesSpace performs spatial clustering at the spot level, utilizing a t-distributed error model to identify clusters, along with employing Markov chain Monte Carlo (MCMC) for estimating model parameters. BASS detects spatial domains and clusters cell types within a tissue section simultaneously by utilizing a hierarchical Bayesian model framework. SpatialPCA is a dimension reduction method aimed at extracting a low-dimensional representation of ST data using spatial correlation information. DR-SC employs a two-layer hierarchical model that simultaneously performs dimension reduction and spatial clustering, optimizing the extraction of low-dimensional features as well as the identification of spatial clusters.

Recent trends indicate a growing momentum toward utilizing graph-based deep learning backbones, attributed to their ability for graphing cell relations and capturing representative features. The representative methods are SpaGCN [21], SEDR [22], CCST [23], STAGATE [3], conST [24], conGI [25], GraphST [4], and ADEPT [26]. These methods predominantly employ graph neural network models to extract latent spot features prior to clustering, albeit with variations in network architectures and design strategies. SpaGCN has a unique design of building an adjacency matrix while considering the histology image pixel values. SEDR employs multiple variation autoen-coders to handle data from different modalities. CCST is based on a graph convolutional network to improve cell clustering and discover novel cell types. STAGATE learns low-dimensional latent embeddings with both spatial information and gene expressions via a graph attention auto-encoder. conST, conGI, and GraphST all emphasize the contrastive learning strategy [27]. conST adopts a two-phase training strategy incorporating self-supervised contrastive learning at three levels: local-local, local-global, and local-context. conGI utilizes three different contrastive learning losses to integrate information from both the histology images as well as the gene expression profiles. GraphST utilizes representations of both normal graphs and corrupted graphs to construct positive and negative spot pairs for contrastive training. ADEPT employs differentially expressed gene selection and imputation procedures to minimize the variations in prediction.

In contrast to merely identifying spatial domains or cell types within a single slice, there is an increasing acknowledgment of the importance of integrative and comparative analyses of multiple ST slices [28]. Thus, ST analysis tools might integrate samples originating from diverse sources, encompassing various individual samples, biological conditions, technological platforms, and developmental stages. Nonetheless, ST slices may exhibit significant “batch effects” [15], which refer to technical biases such as uneven amplification during PCR [29], variations in cell lysis [30], or differences in reverse transcriptase enzyme efficiency during sequencing. These factors have the potential to obscure genuine biological signals, thereby complicating data interpretation and integration.

To analyze multiple ST slices by minimizing batch effects, different alignment and integration methods have been introduced. Three representative alignment methods include PASTE [31], PASTE2 [32], and SPACEL [33]. PASTE utilizes the Gromov-Wasserstein optimal transport (OT) algorithm [34] for aligning adjacent consecutive ST data. PASTE2, an extension of PASTE, allows partial alignment, accommodating partial overlap between aligned slices and/or slice-specific cell types. Both PASTE and PASTE2 output a mapping matrix for every pair of consecutive ST slices, facilitating the reconstruction of the tissue’s 3D architecture through multi-slice alignment. SPACEL combines a multi-layer perceptron and a probabilistic model for deconvolution, subsequently employs a graph convolutional network with adversarial learning to identify spatial domains across multiple ST slices, and finally constructs the 3D tissue architecture by transforming and stacking the spatial coordinate systems of consecutive slices.

Several integration methods have also been introduced. Notable examples include STAligner [35], DeepST [36], PRECAST [37], and SPIRAL [38]. These tools do not directly align slices; instead, they learn shared latent spot embeddings after jointly training on multiple slices. STAligner, built on the STAGATE model, introduces triplet loss by utilizing mutual nearest neighbors between spots from consecutive slices to exploit the contrastive learning strategy for enhancing inter-slice connection. DeepST consists of a graph neural network autoencoder and a denoising autoen-coder to generate a representation of the augmented ST data, as well as domain adversarial neural networks to integrate ST data. PRECAST leverages a unified model including a hidden Markov random field model and a Gaussian mixture model to simultaneously tackle low-dimensional embedding estimation, spatial clustering, and alignment embedding across multiple ST datasets. SPIRAL employs a graph autoencoder backbone with an OT-based discriminator and a classifier to remove the batch effect, align coordinates, and enhance gene expression. BASS also applies the hierarchical Bayesian model framework for multi-slice clustering and outputs clustering labels. The dichotomization of alignment and integration methods is not absolute. PASTE also outputs an integrated center slice, so it can also be classified as an integration tool. STAligner and SPIRAL are also capable of aligning multiple adjacent slices to construct a 3D architecture. For simplicity, we classified each tool into either the alignment or integration category.

Although clustering, alignment, and integration methods have enhanced our understanding of ST data and their practical applications, the lack of comprehensive benchmarking constrains our comprehension and hampers further algorithm development. It is a common phenomenon for a method to demonstrate decent performance on well-studied, commonly used datasets. However, its performance can vary significantly when applied to brand-new data. In this work, we systematically analyze and evaluate the performance of 14 state-of-the-art clustering methods, three alignment methods, and five integration methods on a multitude of simulated and real ST datasets. We design a comprehensive benchmark framework in Fig. 1, and evaluate the clustering performance, overall robustness, layer-wise and spot-to-spot alignment accuracy, integration performance, 3D reconstruction, and computing time of each method. We consolidate these findings into a comprehensive recommendation spanning multiple aspects for the users, while also spotlighting potential areas in need of further research.

**Figure 1.**
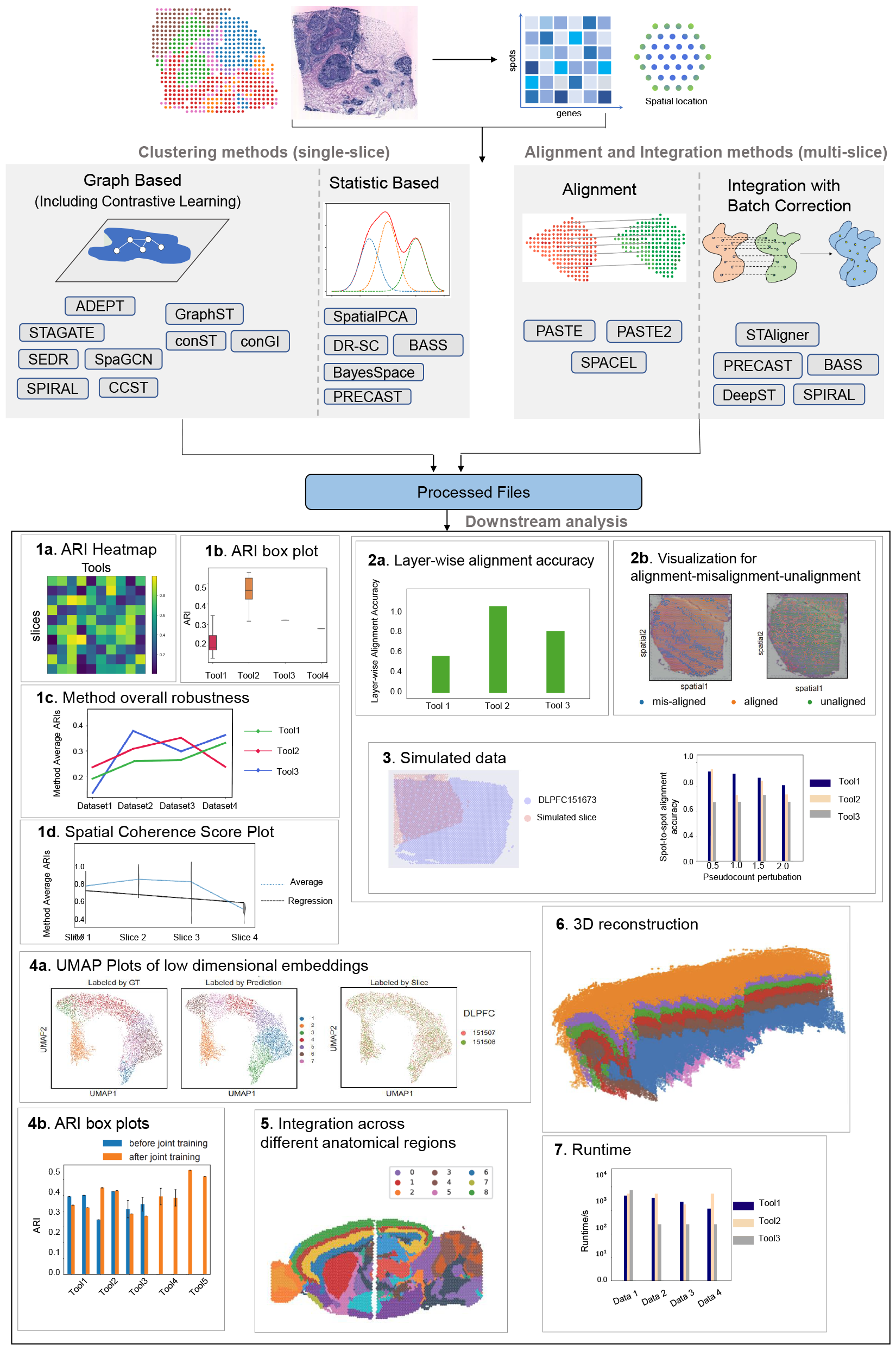
The benchmarking framework for clustering, alignment, and integration methods on different real and simulated datasets. It includes 14 clustering methods, three alignIntegration methods, and five deconvolution methods. Different experimental metrics and analyses, like adjusted rand index (ARI), uniform manifold approximation and projection (UMAP) visualization, layer-wise and spot-to-spot alignment accuracy, spatial coherence score (SCS), and 3D reconstruction, runtime are meticulously designed to quantitatively and qualitatively assess method performance as well as data quality.

## Results

### ST datasets examined and data preprocessing

We collected 10 ST datasets with a total of 69 slices for benchmarking, which had corresponding manual annotations (Table 1). These datasets were produced by several ST protocols, including 10x Visium, ST, Slide-seq v2, Stereo-seq, STARmap, and MERFISH. We broadly categorized them into two groups based on the methodology employed - sequencing-based or imaging-based. The datasets varied in size, withnumber of spots ranging from approximately 1,000 to nearly 10,000, and number of genes from 150 to approximately 36,000.

**Table 1.**
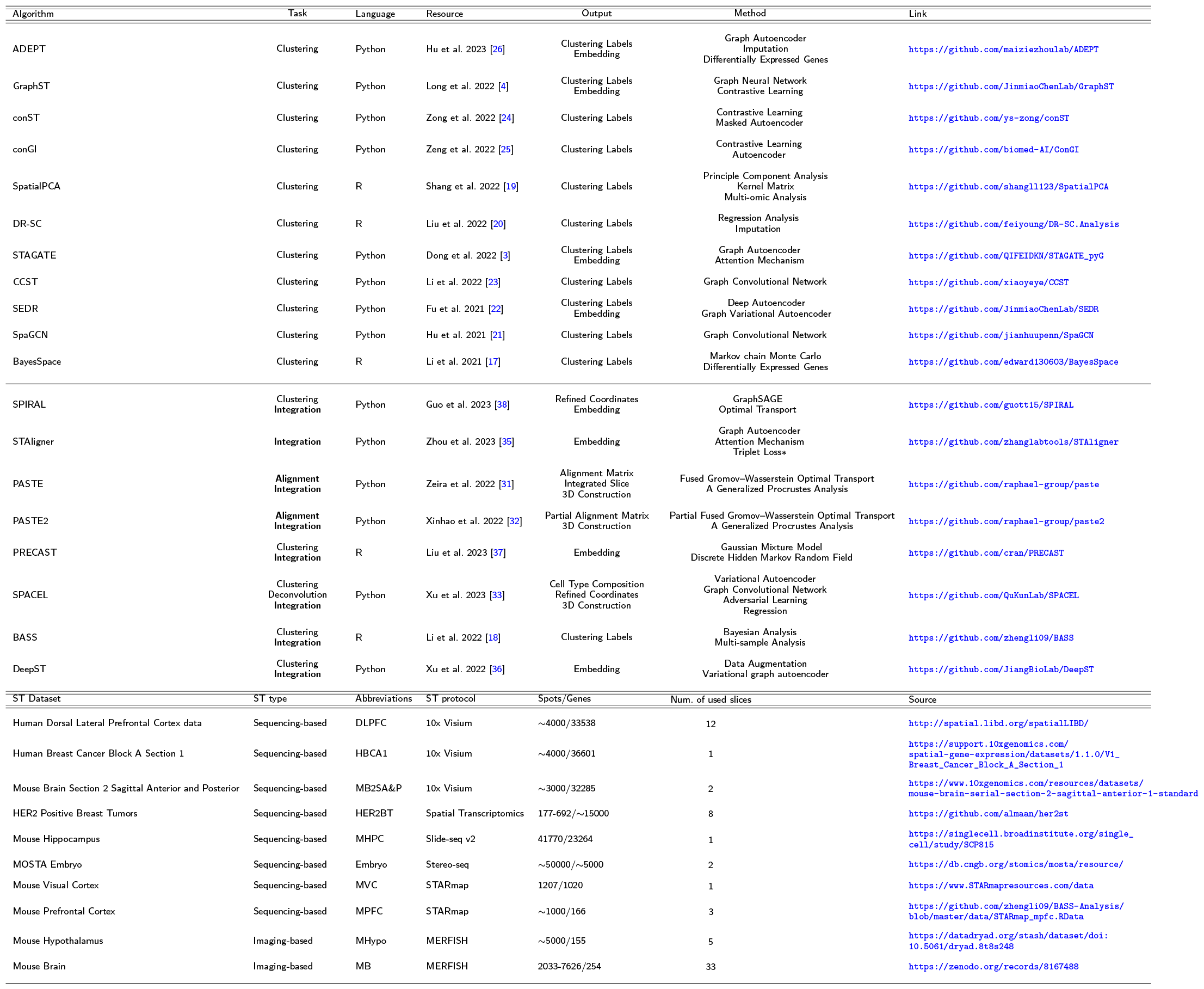
Benchmark tools and datasets. Top panel: Summary of clustering, alignment, and integration methods used in this work. The tool’s task, programming language, resource, tool output, the general method by each tool, and tool links are shown in the table. Bottom panel: Overview of the datasets benchmarked in this study. The ST type, datasets’ abbreviations, ST protocol, number of spots and genes, number of slices, and source link for each dataset are shown in the table.

Specifically, (1) the DLPFC dataset, generated with 10x Visium, includes 12 human DLPFC sections with manual annotation (indicating cortical layers 1 to 6 and white matter (WM)), taken from three individual samples [39]. Each sample contains four consecutive slices.

(2)The HBCA1 dataset, generated with 10x Visium, includes a single slice of human breast cancer, which is open-sourced from 10x genomics [22].

(3)The MB2SA&P dataset, generated with 10x Visium, includes two slices of the anterior and posterior mouse brain. Only the anterior section has annotation [18].

(4)The HER2BT dataset [40] by spatial transcriptomics contains HER2-positive tumors from eight individuals (patient A-H). Each slice has 700 spots and was examined and annotated by a pathologist based on morphology. Regions were labeled as either: cancer in situ, invasive cancer, adipose tissue, immune infiltrate, breast glands, or connective tissue.

(5)The MHPC dataset [41] by Slide-seq v2 is the largest slice used in our study with over 40,000 spots and 23,000 genes. The Allen Mouse Brain Atlas was used as ground truth to identify seven key anatomical regions of the hippocampus, namely CA1, CA2, CA3, dentate gyrus (DG), third ventricle (V3), medial habenula (MH), and lateral habenula (LH).

(6)The Embryo dataset by Stereo-seq has over 50 slices, and the slices at two different time points E11.5 and E12.5 were used in our experiments. These datasets are from a large stereo-seq project called MOSTA [42]: Mouse Organogenesis Spatiotemporal Transcriptomic Atlas by BGI.

(7)The MVC dataset [9] by STARmap contains one slice and was generated from the mouse visual cortex. It extends from the hippocampus (HPC) to the corpus callosum (CC), and includes the six neocortical layers.

(8)The MPFC dataset [9] of the mouse prefrontal cortex, annotated by BASS [18], was sequenced with the STARmap protocol with expression values measured for 166 genes on 1127 single cells, along with their centroid coordinates on the tissue. Spatial domains, such as cortical layers L1, L2/3, L5, and L6, have been assigned based on the spatial expression patterns of marker genes, including Bgn for L1, Cux2 for L2/3, Tcerg1l for L5, and Pcp4 for L6. Three slices in this dataset are not categorized as consecutive.

(9)The MHypo dataset by MERFISH contains five manually annotated consecutive slices [18] labeled Bregma-0.04 (5488 cells), Bregma-0.09 (5557 cells), Bregma-0.14 (5926 cells), Bregma-0.19 (5803 cells), and Bregma-0.24 (5543 cells). Expression measurements were taken for a common set of 155 genes. Each tissue slice includes a detailed cell annotation, identifying eight structures: third ventricle (V3), bed nuclei of the stria terminalis (BST), columns of the fornix (fx), medial preoptic area (MPA), medial preoptic nucleus (MPN), periventricular hypothalamic nucleus (PV), paraventricular hypothalamic nucleus (PVH), and paraventricular nucleus of the thalamus (PVT).

Finally, (10) the MB dataset [33] by MERFISH has 33 consecutive mouse cerebral cortex tissue slices with similar shapes, which were used for 3D reconstruction. Annotation includes the six layers (L1-L6) and distinct neuronal cell types such as intratelencephalic (IT), sub-cerebral projection, and cortico-thalamic (CT) neurons.

All except the Embryo and MB datasets were used for benchmarking clustering tools. Five datasets, DLPFC, MB2SA&P, Embryo, MHypo, and MB, were used for benchmarking alignment and integration tools.

Most methods have uniquely designed procedures for data preprocessing. The preprocessing step could significantly impact the sensitivity of these methods. We provided the specific pipeline of data preprocessing for each method in our GitHub. In this context, we concentrated on the common overlapping steps employed by the majority of methods. The preprocessing of ST data typically encompasses three essential steps: removal of extreme values, normalization, and dimension reduction. The scanpy package is often utilized to eliminate low-quality cells lacking sufficient expressed transcripts or low-quality genes rarely observed across the data slice, thus mitigating the impact of noise. Subsequently, the expression matrix is normalized within each cell and log-transformed to further suppress potential extreme values. Finally, if the feature selection step is enabled, dimension reduction can be achieved through methods such as highly variable gene selection or principal component analysis (PCA). This enables models to focus on essential features rather than high-dimensional raw inputs.

Utilizing the evaluation framework illustrated in Fig. 1, we conducted benchmarking of various clustering, alignment, and integration methods across all ST datasets.

### Performance comparison of 14 clustering methods

We first performed a comprehensive benchmarking analysis for 14 different clustering methods aimed at assessing their performance in accurately identifying spatial domains or cell types. The two heatmaps of Fig. 2a-b illustrated the average Adjusted Rand Index (ARI) for each method across 31 slices from seven ST datasets, along with the corresponding rank scores for each tool. The detailed method for computing ARI values and each rank score is outlined in the methods section. The ARI and rank results revealed that STAGATE, GraphST, ADEPT, and BASS emerged as top-tier tools. Notably, BASS attained the highest sum and average rank, followed by GraphST and ADEPT. BASS achieved a much higher ARI than other methods on the MHypo datasets. Most methods struggled to give reasonable predictions on the HER2BT datasets since the annotated regions by ground truth were less coherent. This comprehensive evaluation shed light on the relative strengths of these methods in the context of spatial domain identification within each ST slice. In Fig. 2j, we further present a holistic assessment of the overall robustness of each clustering method by aggregating the average ARI across slices within each of the seven datasets and depict the results in a line chart. Notably, lower variances were exhibited in the DLPFC, HER2BT, and the anterior section of MB2SA&P datasets across all clustering methods, albeit for different reasons. BASS, in alignment with previous analyses, emerged as the best clustering tool for four datasets. Nevertheless, it exhibited comparatively poorer performance on the HBCA1 dataset. GraphST and ADEPT consistently secured the second and third positions, respectively, across most datasets, with GraphST taking the lead in the DLPFC and the anterior section of MB2SA&P datasets. This evaluation offers insight into the performance of various methods across diverse datasets, further highlighting both strengths and challenges.

**Figure 2.**
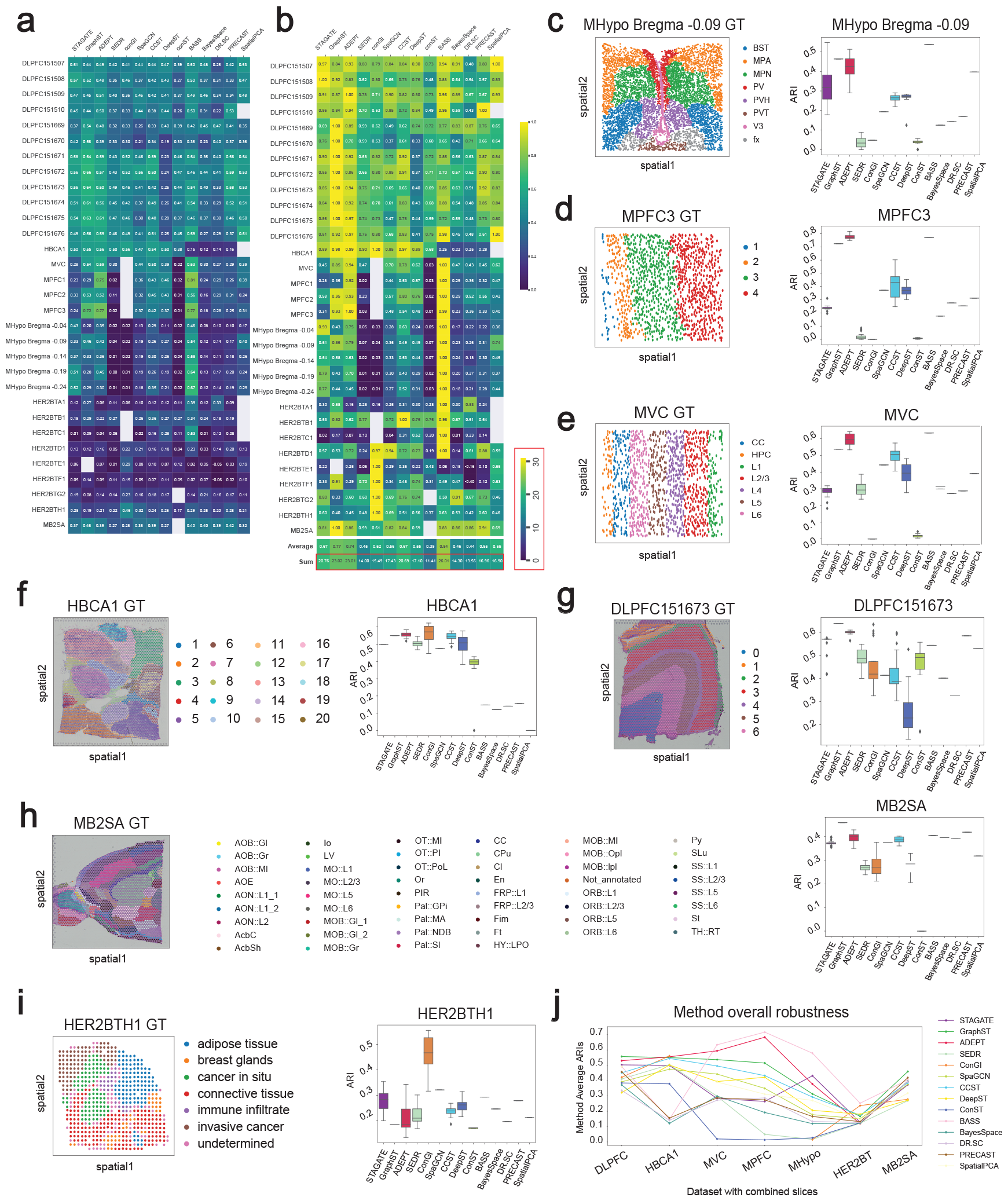
Clustering performance over 14 methods on 31 ST slices of seven datasets. **a** ARI heatmap. Each average ARI value was based on 20 runs. **b** Ranking heatmap. This ranking heatmap was created by normalizing all results within the same slice by dividing them by the maximum ARI value (representing the best performance) among all methods, thus standardizing all ARI values to 1. For each method, the best ranking for the sum result is 31, and the best ranking for the average result is 1. **c-i** The ground truth visualization plots and box plots depicting ARI values of all tools on selected data slice from each dataset. **j** line plots displaying the overall robustness of all methods across seven datasets.

Since the mean ARI failed to capture the variance of each method, we also plotted boxplots and ground truth visualization plots on all slices from each dataset (In Fig. 2c-i and Fig. S1-S2). Five statistical methods, namely BASS, BayesSpace, DR.SC, PRECAST, and SpatialPCA, exhibited no variance as they yielded deterministic outputs. The remaining methods primarily relied on graph-based deep learning techniques, leading to potential variations in their predictions owing to random seeds. However, GraphST, ConGI, and SpaGCN fixed their seeds to be identical for each run. So far, we quantitatively evaluated all clustering methods by ARI. For the MHPC data (Fig. 3a), where the spots were labeled by cell types, visual comparison with the ground truth was more effective than calculating ARIs. Additionally, we employed the Allen Brain Atlas as a ground truth for the anatomical regions. The ground truth comprised four key distinguished anatomical regions, CA1, CA2, CA3, and dentate gyrus (DG), which displayed curved shapes. Our results demonstrated that all methods successfully recovered this feature, however, DR.SC and BASS failed to identify them as separate regions. Moreover, ADEPT, GraphST, and STAGATE could further differentiate CA1 and CA3. Notably, no method delineated a separate CA2 region, merging it with CA3 instead. We further investigated three other key anatomical regions - third ventricle (V3), medial habenula (MH), and lateral habenula (LH) - and only BASS, ADEPT, and STAGATE could successfully delineate these three regions. In conclusion, ADEPT and STA-GATE were the best tools for delineating all seven key regions.

**Figure 3.**
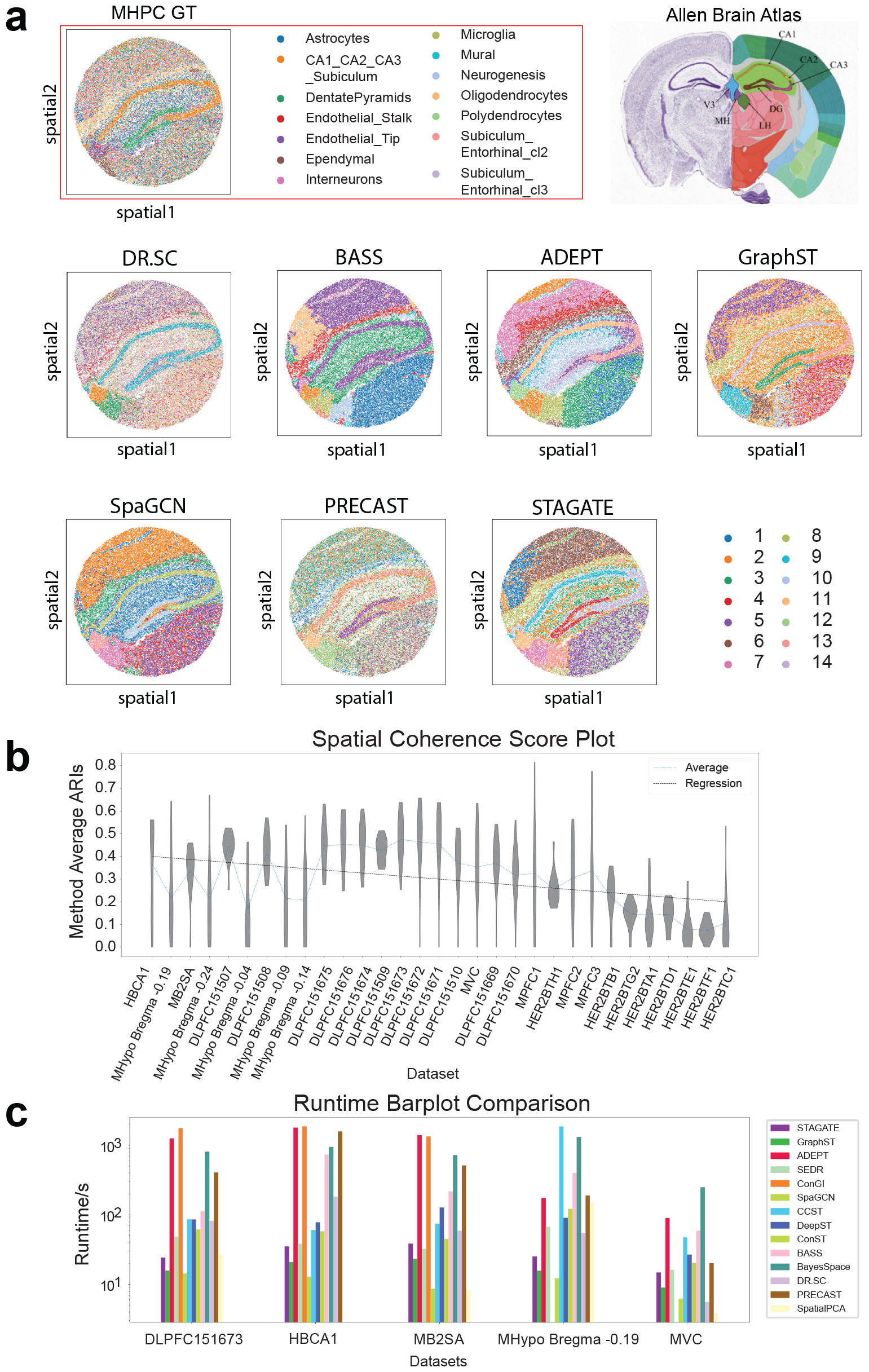
Clustering visualization on the MHPC dataset, data complexity tendency using spatial coherence score, and runtime comparison of all clustering methods. **a** Comparisons of the predicted clusters generated by different clustering methods with the ground truth for the MHPC data. **b** Violin plots with the regression line showing the decline in average ARI across all methods as data complexity increases across 31 ST slices. Using spatial coherence score as an indicator of data complexity. **c** Runtime analysis of all clustering methods.

### Spatial coherence score and runtime analysis for clustering methods

All clustering methods exhibited varied performance across datasets. To reveal the effect of data complexity on performance, we reordered all 31 data slices based on their complexity from low to high and demonstrated the violin plots of all methods’ performance on each data along with the regression and mean value line plots (Fig. 3b and Fig. S3). The Spatial Coherence Score (SCS) was introduced as a metric for quantifying data complexity. The detailed method is described in the methods section. The overall tendency of the average ARI across all methods, represented by the regression line, decreased as the data complexity increased. However, there were some fluctuations due to other factors. An intriguing observation emerged: although a discernible overall trend was evident, the average ARIs on well-studied datasets were mostly above the regression line, whereas for less-studied datasets, average ARIs lagged below the regression line. This outcome indicated that the designs of most current algorithms favored the commonly used datasets and were not generally effective for all datasets. Though this phenomenon was due to the scarcity of available ST datasets with high-quality ground truth, it did exhibit a potential issue of algorithm overfitting, which should be noted and prevented in future studies.

Finally, we benchmarked the runtime of each method on five selected datasets (Fig. 3c). Three of them, the DLPFC 151673, HBCA1, and the anterior section of MB2SA&P datasets, are large with approximately 4k spots and 20-30k genes. Though each slice of the MHypo dataset has approximately 5k spots, each spot only contains 155 genes. The MVC dataset is the smallest in terms of the number of spots and genes. The runtime of each method was similar across the first three datasets due to the size similarity of the data. Overall, STAGATE, GraphST, and SpaGCN, exhibited dominant advantages in terms of runtime, based on the fact that they could analyze most of the data slice within a minute. SEDR, CCST, DeepST, ConST, BASS, and DR.SC had comparably worse speed, while they could still finish execution in about 5 minutes. ADEPT, ConGI, BayesSpace, and PRECAST lacked scalability and were greatly affected by both the number of spots and genes since their time consumption increased drastically when the data size increased. In contrast, methods like GraphST and STAGATE had better scalability and were more resistant to the data size by maintaining a general uniform time consumption.

### Assessing the characteristics of joint spot embedding with pairwise two-slice joint analysis

In contrast to the conventional approach of ST focusing on spatial domain distribution in a single slice, there is a growing recognition of the value of integrative and comparative analyses of ST datasets. In our pairwise two-slice joint analysis, we started by using nine pairs of DLPFC slices to explore whether integration could improve joint spot embeddings by leveraging adjacent consecutive slices. Evaluation experiments were conducted by introducing layer-wise alignment accuracy. The fundamental idea behind this analysis is based on the hypothesis that aligned spots across consecutive slices are more likely to belong to the same spatial domain or cell type. The detailed method for defining layer-wise alignment accuracy is outlined in the methods section.

In Fig. 4a, we compared the layer-wise alignment accuracy of all seven methods on nine DLPFC slice pairs. Given the unique layered structure of DLPFC data, we designed this evaluation metric to assess whether “anchor” spots from the first slice and “aligned” spots from the second slice belong to the same layer (layer shifting = 0) or different layers (layer shifting = 1 to 6). The expectation was that a good integration or alignment tool would show high accuracy for anchor and aligned spots belonging to the same layer (layer shifting = 0), and this accuracy should decrease when the number of layer shifting increases. In seven out of nine DLPFC slice pairs, SPACEL demonstrated the highest layer-wise alignment accuracy, while PASTE led in the remaining two pairs (Fig. 4a). A similar experiment was conducted on four pairs drawn from the MHypo dataset (Fig. 4b), SPACEL still exhibited the best performance, followed by PASTE in the second position. It was not surprising that the two alignment tools, SPACEL and PASTE, exhibited the highest accuracy in layer-wise alignment, which was expected as their primary objective was the direct alignment of spots across slices, rather than relying on joint spot embeddings for integration analysis. Conversely, tools like STAligner, PRECAST, DeepST, and SPIRAL, which leverage joint spot embeddings for indirect alignment across slices, demonstrated slightly lower but still satisfactory layer-wise alignment accuracy. Among these tools, STAligner achieved the highest accuracy, followed by DeepST, while PRECAST performed the least accurately. These results highlighted, to some extent, the inherent qualities of joint spot embeddings by these integration tools. PASTE2, an extension version of PASTE, exhibited poor performance in this scenario because it primarily addresses the partial overlap alignment problem, where only partial overlap occurs between two slices or slice-specific cell types.

**Figure 4.**
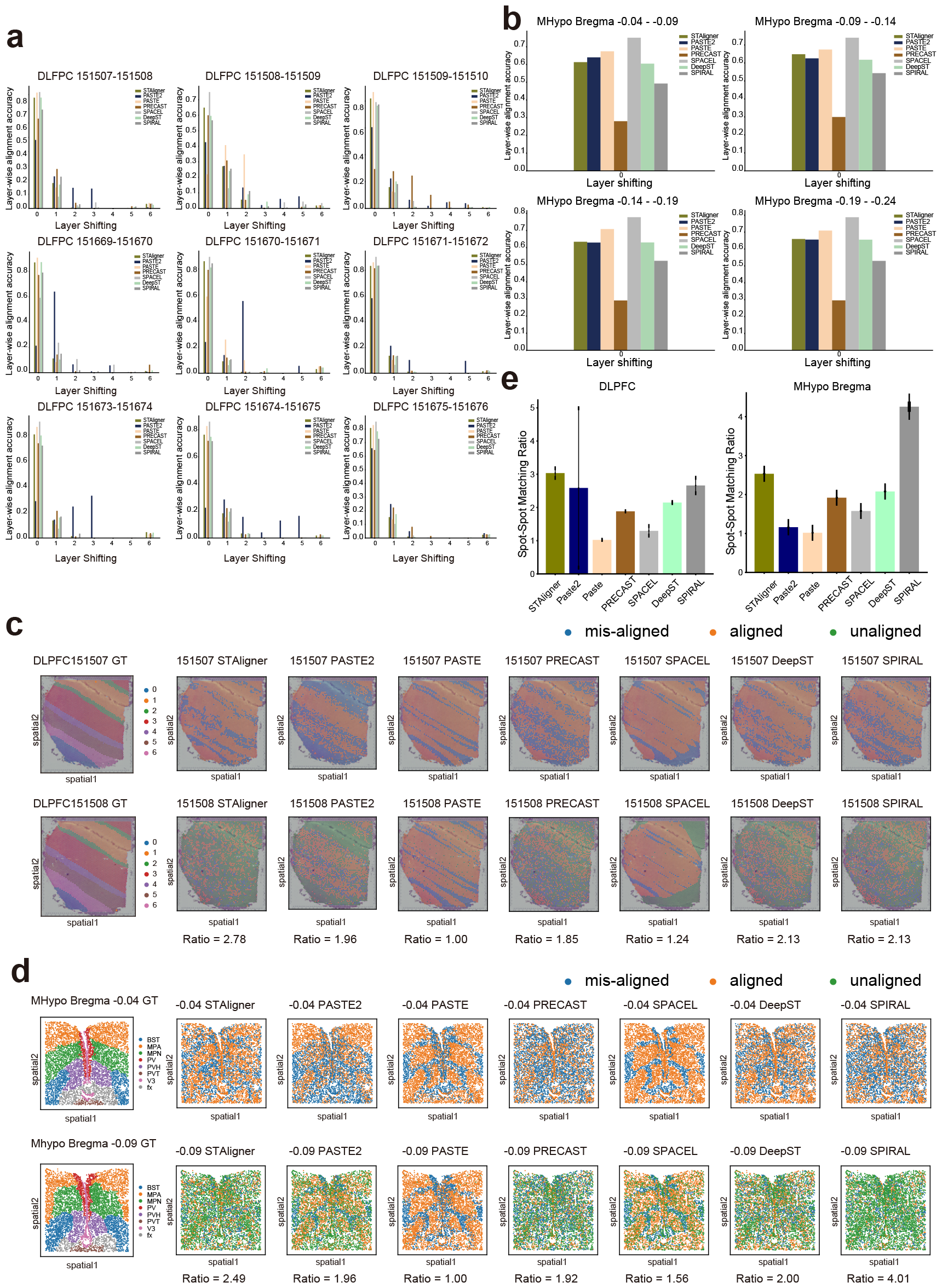
Bar plots for layer-wise alignment accuracy and spot-to-spot mapping ratio, and visualization plots for alignment-misalignment-unalignment. **a-b** Bar plots depicting the layer-wise alignment accuracy of seven methods on nine DLPFC slices (a) and four MHypo slices (b). **c-d** Visualization plots showing aligned spots, misaligned spots, and unaligned spots when aligning the anchor spot from the first (top) slice to the aligned spots on the second (bottom) slice on DLPFC 151507-151508 and MHypo Bregma -0.04 - -0.09 pair. Values below each plot represent the spot-to-spot matching ratio. **e** Bar plots representing the average spot-to-spot mapping ratio of each tool on two datasets: DLPFC and MHypo.

While layer-wise alignment accuracy provides insight into spot-to-layer alignment, it is crucial to evaluate the spot-to-spot matching ratio to further evaluate joint spot embeddings. In Fig. 4c-d, we marked “anchor” and “aligned” spots on both slices using three different colors, further classifying them into aligned (orange), misaligned (blue), and unaligned (green) spots based on ground truth layer labels, as described in the methods section. Notably, for the DLPFC 151507 - 151508 pair, STAligner, PASTE2, PRECAST, DeepST, and SPIRAL showed a notable proportion of unaligned spots on the second slice. This suggested a bias in these five tools, aligning multiple “anchor” spots from the first slice to the same “aligned” spot on the second slice, thereby leaving a significant number of spots unaligned on the second slice. The spot-to-spot mapping ratio further corroborated this observation, with PASTE demonstrating the lowest ratio (1.00), followed by SPACEL (1.24), PRE-CAST (1.85), PASTE2 (1.96), DeepST (2.13), SPIRAL (2.13), and STAligner (2.78). Averaging this ratio across all nine pairs for each tool revealed a consistent pattern (Fig. 4e). Moreover, across all nine pairs, it was observed that misaligned spots (Fig. 4c and Fig. S4-S5) on the first slice tended to aggregate along the layer boundaries in PASTE and SPACEL. In contrast, other tools such as STAligner, PASTE2, PRECAST, DeepST, and SPIRAL, exhibited a dispersion of these misaligned spots within the layers. Specifically, PRE-CAST displayed the most dispersed pattern, which was in agreement with its least favorable layer-wise alignment accuracy. The low spot-to-spot mapping ratio and the dispersed pattern of misaligned spots in all integration tools suggested a shared trade-off, wherein the learned low-dimensional embeddings sacrifice certain local geometric information in the process of optimization and training. SPACEL, the alignment tool, exhibited coherent regions of unaligned spots (illustrated in green) outside the matched regions.

We further performed this evaluation analysis in four pairs of MHypo slices and observed a similar trend for spot-to-spot mapping ratio and a similar dispersed pattern of misaligned spots in all tools (Fig. 4d and Fig. S6). Specifically, SPIRAL had the worst average spot-to-spot mapping ratio, followed by STAligner, DeepST, and PRECAST (Fig. 4e). PASTE2 and PASTE achieved a ratio of approximately 1. SPACEL demonstrated a less favorable average ratio (1.58) for the MHypo data in comparison to the DLPFC data (1.30).

### Performance of alignment accuracy on simulated datasets

While real datasets enabled us to assess alignment accuracy to some extent, they lacked precise spot-to-spot alignment ground truth. To comprehensively investigate alignment accuracy, we simulated datasets with the gold standard for different scenarios to demonstrate the robustness of all alignment and integration methods.

We first used one DLPFC slice as the reference and simulated another slice with different overlap ratios (20%, 40%, 60%, 80%, and 100%) in comparison to the reference slice (Fig. 5a). In this simulation scenario, the pseudocount (gene expression) perturbation was fixed at 1.0 for all simulated slices. The detailed simulation method is outlined in the methods section. In Fig. 5b (left panels), the layer-wise alignment accuracy and spot-to-spot alignment accuracy were shown in the bar plots. We observed that the layer-wise alignment accuracy of all methods tended to decrease when the overlapping ratio between two slices decreased. Nevertheless, in terms of spot-to-spot alignment accuracy, all four integration methods - STAligner, PRECAST, DeepST, and SPIRAL - failed to achieve even a marginal value, which was consistent with the earlier conclusion that these tools exhibit relatively high spot-to-spot mapping ratios. On the other hand, three alignment tools - SPACEL, PASTE2, and PASTE - achieved relatively better spot-to-spot alignment accuracy. Among them, PASTE2 achieved a near-perfect accuracy at the 100% overlapping ratio and consistently maintained approximately 60% accuracy at lower overlapping ratios. SPACEL exhibited slightly better accuracy than PASTE2 when the overlapping ratio was lower than 100%. However, its accuracy decreased to approximately 40% at the 100% overlapping ratio. PASTE, on the other hand, failed to achieve satisfactory accuracy when the overlapping ratio was lower than 100%.

**Figure 5.**
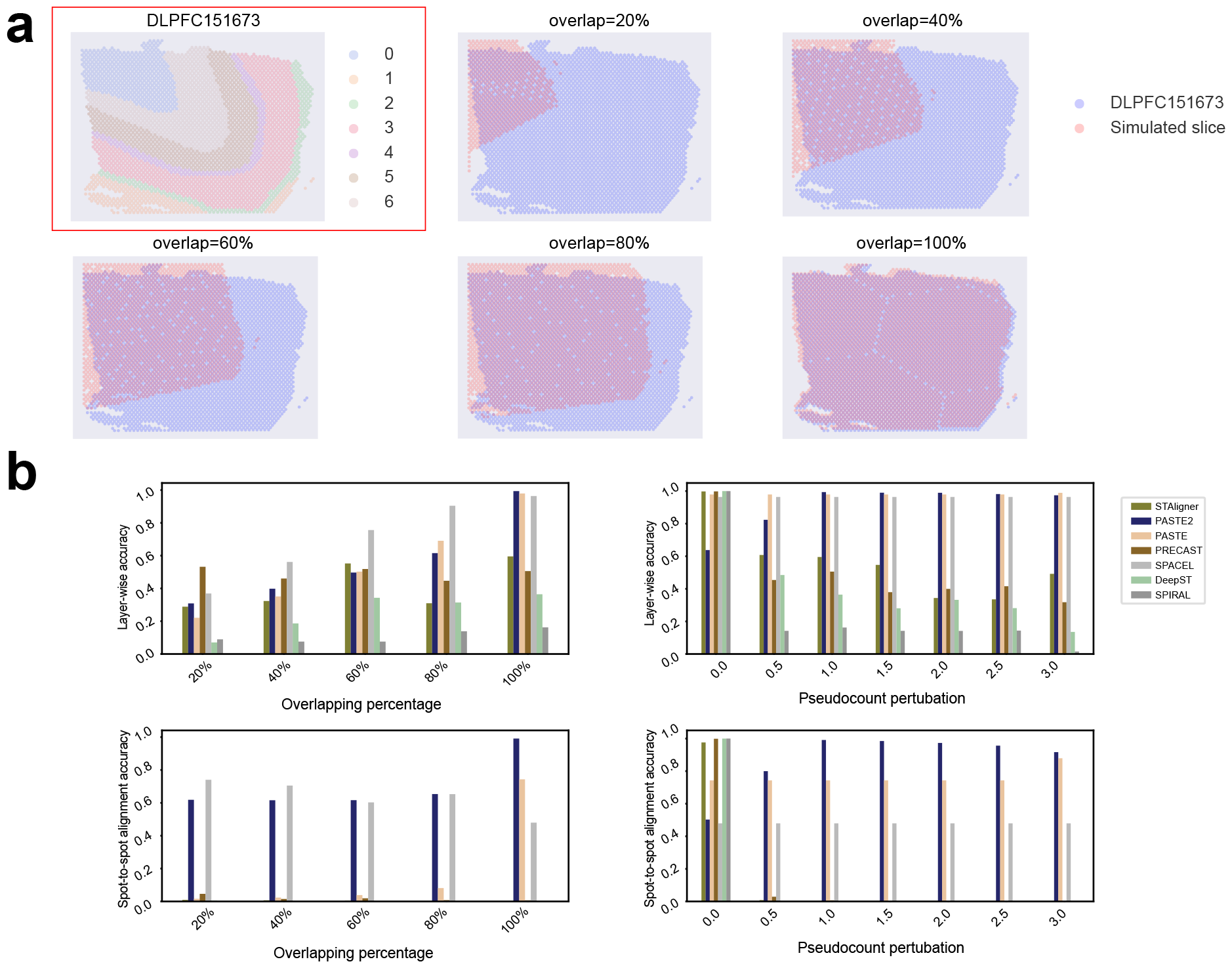
Alignment accuracy in simulation Data. **a** DLPFC 151673 slice, consisting of seven layers, along with its simulated consecutive slices featuring overlapping ratios of 20%, 40%, 60%, and 80% with respect to DLPFC 151673 slice. **b** Layer-wise accuracy and spot-to-spot accuracy of different tools when overlapping ratio increases or when pseudocount perturbation increases.

In the second simulation scenario, we simulated the slice with different pseudocounts (0 - 3.0 with a step size of 0.5) to represent perturbation on gene expression while keeping the overlapping ratio fixed at 80%. In Fig. 5b (right panels), the bar plots demonstrated that the layer-wise alignment accuracy of four integration tools - STAligner, PRECAST, DeepST, and SPIRAL - decreased when pseudocount perturbation increased. This result suggested that all integration methods were sensitive to perturbation on the expression profiles, as they utilized gene expression profiles as spot (node) features when constructing a graph model for training. Conversely, three alignment tools - SPACEL, PASTE2, and PASTE - exhibited significantly greater resilience to perturbations in gene expression. This resilience stems from their objective functions for alignment, which allowed for a more pronounced emphasis on spatial coordinates when gene expression varied across slices. Regarding spot-to-spot alignment accuracy, these three alignment tools consistently maintained similar accuracy across various pseudocount perturbations. PASTE2 demonstrated the highest accuracy when pseudocount perturbation ranged from 0.5 to 3.0. Notably, when pseudocount perturbation was set to 0, indicating identical gene expression levels for each spot across slices, all four integration tools achieved better accuracy.

### Integration methods improve integration of consecutive slices with batch correction

Once joint spot embeddings for each integration method were generated, we further visually evaluated the “batch-corrected” joint embeddings for the integration of consecutive slices using two components from UMAP. Alignment tools, PASTE, PASTE2, and SPACEL, were excluded from this analysis as they did not generate latent embeddings. For the DLPFC 151507 and 151508 pair (Fig. 6a), the UMAP plots for PRECAST, STAligner, DeepST, and SPIRAL showed that spots from two different slices were evenly mixed to some extent (Fig. 6a, right panel), and their predicted domain clusters were well segregated (Fig. 6, middle panel). Specifically, PRE-CAST tended to generate embeddings in a pattern with separated clusters, with some predicted clusters encompassing spots from different domains, a pattern that did not entirely align with the ground truth (Fig. 6a, left panel). STAligner, DeepST, and SPIRAL maintained the hierarchical connections of the seven layers in the latent embedding space to some degree. However, there were instances where predicted spatial domains included spots from nearby domains, or one spatial domain was predicted to be two adjacent domains. STAligner achieved better UMAP visualization than DeepST and SPIRAL. Among all tools, PRECAST lost more geometry information than the other three tools since it prominently separated spatial domains in the latent space. We further demonstrated this UMAP analysis for all the rest DLPFC pairs and plotted annotations by ground truth, method prediction, and slice index (Fig. S7-S8). All remaining UMAP results exhibited consistent patterns and further affirmed that all four methods were capable of generating “batch-corrected” joint embeddings for the integration of consecutive slices. However, the integrated spatial domains were not highly concordant with the ground truth.

**Figure 6.**
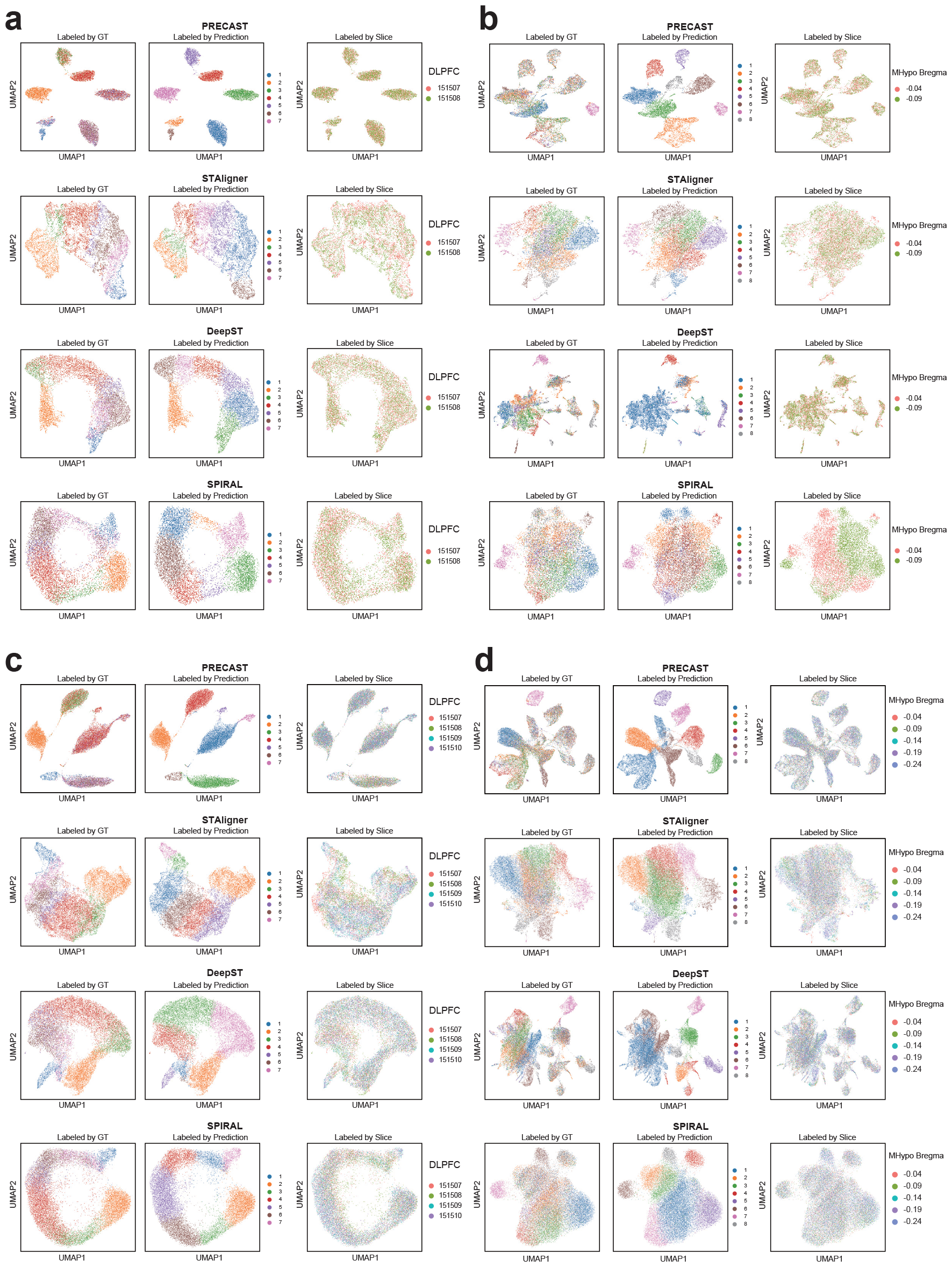
UMAP plots of low dimensional joint embedding distribution for batch correction. **(a-d)**These UMAP plots depicting the 2D distribution of latent joint embeddings after integration with batch correction by different methods on the DLPFC 151507-151508 pair (a), the MHypo Bregma -0.04 - -0.09 pair (b), the DLPFC 151507-151510 four slices (c), and the MHypo Bregma -0.04 - -0.24 four slices. Each UMAP contains colored spots labeled by three different setups: ground truth (GT), method prediction, and slice index.

We extended this analysis to four pairs of the MHypo data (Fig. 6b and Fig. S9). The joint embeddings generated by PRECAST, STAligner, and DeepST somewhat facilitated integration across consecutive slices, although this effect was much inferior compared to the results of the DLPFC data. These three tools exhibited several connected small clusters or a single large cluster which were hard to differentiate based on the annotation by ground truth. The other tool, SPIRAL, experienced a significant batch effect as its joint embeddings across slices were unevenly mixed and experienced substantial separation. This result was in agreement with the least favorable spot-to-spot mapping ratio (4.25) by SPRIAL.

In addition to benchmarking on the integration of slice pairs, we further demonstrated the performance of each method on multi-slice (*>* 2) integration. All UMAP plots for PRECAST, STAligner, DeepST, and SPIRAL indicated a relatively even mixture of spots from four distinct slices provided by three samples (DLPFC 151507-151510, 151669-151672, 151673-151676) (Fig. 6c and Fig. S10). Consistent with observations in paired settings, the embeddings generated by PRECAST continued to exhibit a pattern characterized by separated clusters. On the other hand, STAligner, DeepST, and SPIRAL still maintained hierarchical connections across seven layers in the latent embedding space. STAligner demonstrated slightly better UMAP visualization than DeepST and SPIRAL. As for the integration of the five slices of the MHypo dataset (Fig. 6d), all tools still displayed several small connected clusters or a single large cluster that was challenging to differentiate based on the annotation by ground truth. However, SPIRAL mixed the spots across five slices evenly and did not display any batch effect, which indicated SPIRAL could use adequate data to remove the batch effect for its latent embeddings. In summary, there is still a need for an optimal and robust tool for integration. While existing tools have shown efficacy to some extent in well-studied datasets, their performance has not consistently generalized to diverse datasets.

### Integration methods enhance domain identification through joint embedding

Integrating data from multiple ST slices can allow us to estimate joint embeddings of expressions representing variations between cell or domain types across slices, which has the potential to better detect spatial domains or cell types, compared to single slice analysis [31]. To further quantitatively compare the effectiveness of these methods in capturing spatial domains via joint embeddings, we employed joint embeddings from each pair of slices in the MHypo and DLPFC datasets to perform clustering together using the clustering method mclust [43]. We then computed ARI as an evaluation metric to compare the clustering results of each tool with the ground truth in each slice, with higher ARI scores indicating better domain identification.

In Fig. 7, we plotted the average ARI results under two scenarios. BASS, PRECAST, and DeepST supported both single-slice and multi-slice joint (integration) analyses. Accordingly, we utilized blue bars to depict the results before joint clustering (single-slice mode) and orange bars to represent the results after joint clustering. However, since STAligner and SPIRAL only have a multi-slice joint analysis mode, the blue bars for these methods were left unpopulated.PRECAST and DeepST achieved better clustering results in terms of ARI for most slices after joint clustering. However, it was difficult to conclude which tool had the overall best performance in all pairs after joint clustering. In nine pairs of DLPFC data (Fig. 7a), DeepST and STAligner exhibited the most variance across all runs. SPIRAL demonstrated the best performance on DLPFC 151509 - 151510 and 151669 - 151670 pairs. STAligner led the performance on DLPFC 151673 - 151674, 151674 - 151675, and 151675 - 151676 pairs, albeit marginally. Notably, the DLPFC 151670 - 151671 pair, characterized by a large spatial distance along the z-axis within the tissue between two slices, presented challenges for all methods. These tools either exhibited a significant performance discrepancy in two slices or failed to perform well in both slices. A similar observation has been spotted on the 151508 - 151509 pair as well. In the DLPFC 151671 - 151672 pair, SPIRAL and STAlinger demonstrated the better performance. Most methods performed similarly on the DLPFC 151507 - 151508 pair. Results were comparatively simpler on the four pairs of the MHypo dataset (Fig. 7b). BASS demonstrated superior performance in all four pairs, followed by STAligner. However, the remaining three methods failed to produce reasonable results.

**Figure 7.**
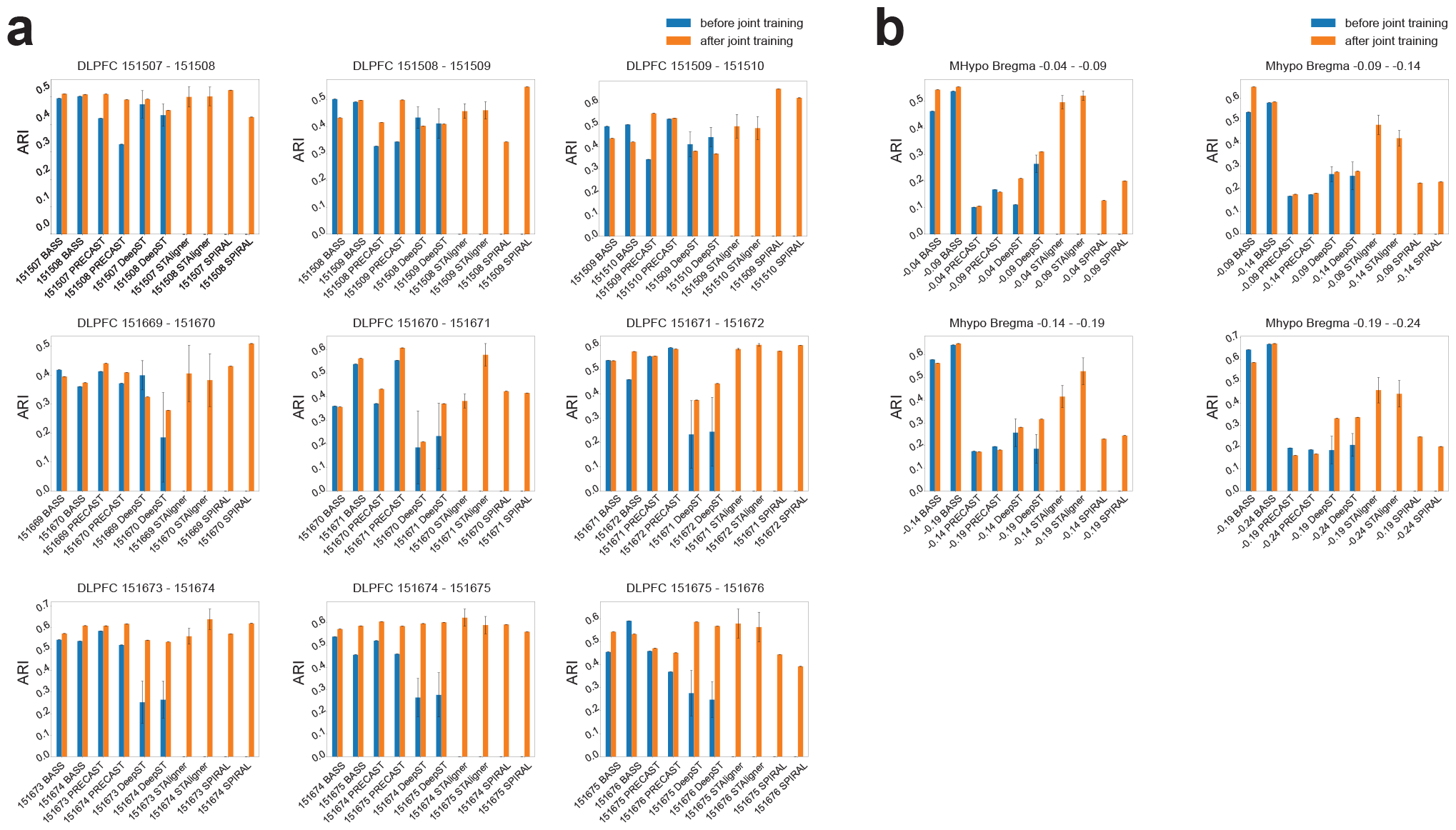
ARI boxplots before and after joint training for domain identification in DLPFC and MHypo datasets. **(a)** ARI boxplots for nine DLPFC pairs by different methods. **(b)** ARI boxplots for four MHypo pairs by different methods. Blue bars represent ARI values before joint clustering (in single-slice mode) and orange bars represent ARI values after joint clustering. The blue bars for STAligner and SPIRAL remain unpopulated since they do not support single-slice clustering.

### Integration methods align samples across different anatomical regions and development stages

So far, our benchmarking has focused on evaluating the integration capabilities of methods across adjacent consecutive sample slices. In this section, we delved deeper into its efficacy for integrating non-consecutive slices. We employed a 10x Visium dataset representing mouse brain sagittal sections, divided into posterior and anterior. We employed the Allen Brain Atlas as a reference (Fig. 8a) and visually compared the clustering results of all methods (Fig. 8b-f). Among all methods, PRECAST demonstrated the least effective performance and failed to detect and connect common spatial domains. In contrast, BASS, STAligner, DeepST, and SPIRAL were better able to identify and connect common spatial domains along this shared boundary. Specifically, only STAligner identified and aligned six distinct layers in the cerebral cortex (CTX) across the anterior and posterior sections. On the other hand, BASS and SPRIAL only managed to identify four distinct layers in CTX. Additionally, STAligner and SPRIAL performed well in distinguishing layers within the cerebellar cortex (CBX). However, none of them identified a coherent arc across two sections for CA1, CA2, and CA3. In summary, STAligner showed capacity in integration for adjacent slices across different anatomical regions.

**Figure 8.**
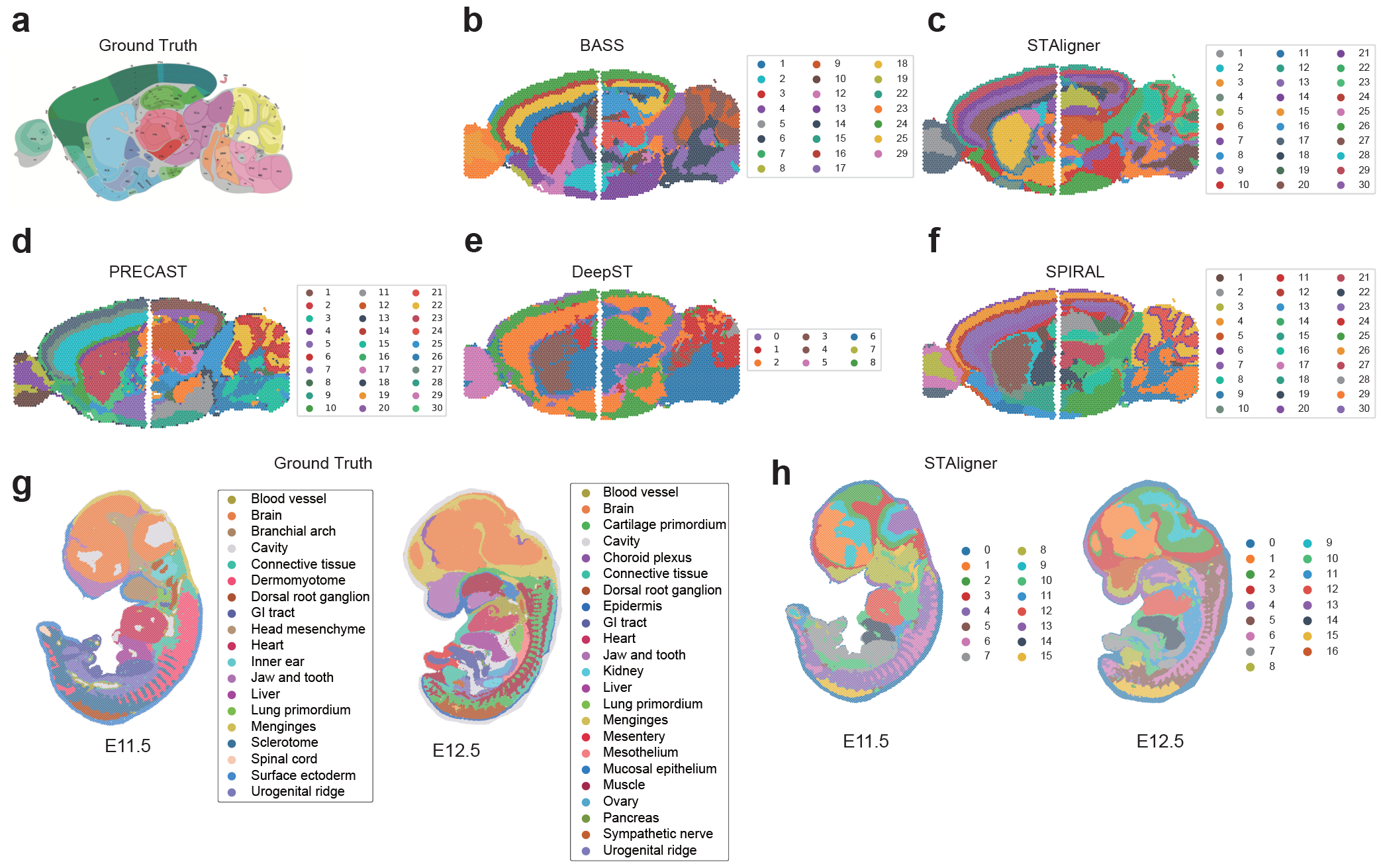
Visualization plots for integration with batch correction in MB2SA&P dataset and mouse Embryo dataset. **a** The Allen Brain atlas serving as the ground truth. **b-f** Domain identification by five methods in the MB2SA&P dataset. **g** Domain identification by the ground truth in the mouse Embryo dataset. **h** Domain identification by STAligner in the mouse Embryo dataset.

Next, we investigated the ability of all methods to integrate two slices from different development stages, to study the spatiotemporal development in tissue structures during mouse organogenesis. Only STAligner has scalability in processing this big benchmarking dataset (over 50k spots for each slice), so other tools were excluded from this analysis. In Fig. 8g, the two mouse embryo slices were acquired at two different time points (E11.5 and E12.5) with region-based manual annotations for different organs and tissues. We observed that STAligner successfully retrieved several shared structures such as dorsal root ganglion, brain, heart, and liver in both slices (Fig. 8h). We also observed that at developmental stage E11.5, structures like the ovary and kidney were less developed compared to E12.5. These results facilitated the reconstruction of the developmental progression of each tissue structure throughout organogenesis.

### Reconstruction of 3D architecture from consecutive 2D slices

Initially, 2D slices were produced from 3D tissue, and alignment tools, specifically designed for pair-wise or all-to-all alignments using multiple adjacent consecutive slices, can then reconstruct the 3D architecture. 3D architecture allows users to explore the dynamics of transcript distributions from any direction, so reconstructing an effective 3D architecture of complex tissues or organs is essential. In Fig. 9, we provided 3D reconstruction visualization results from three different samples using four methods, SPACEL, PASTE, SPIRAL, and STAligner. All four tools achieved consistent and satisfactory 3D visualization results on DLPFC sample 3, encompassing four adjacent consecutive slices numbered 151673-151674-151675-151676 (Fig. 9a). For the MHypo sample which contains five consecutive slices, SPACEL and PASTE demonstrated comparable and effective 3D visualizations (Fig. 9b). In contrast, SPIRAL exhibited mis-aligned scatter spots beginning from the second slice, and the occurrence of these misalignments increased with the addition of more stacks of slices. Starting from the third slice, STAligner exhibited rotational distortions in the slices, leading to a discordant 3D architecture. The underlying reason could be that SPIRAL performed all-to-all alignments, whereas SPACEL and PASTE performed pairwise alignments between each pair of adjacent consecutive slices sequentially. All-to-all alignments have the potential to introduce more false alignment, particularly when two slices are not closely positioned along the z-axis.

**Figure 9.**
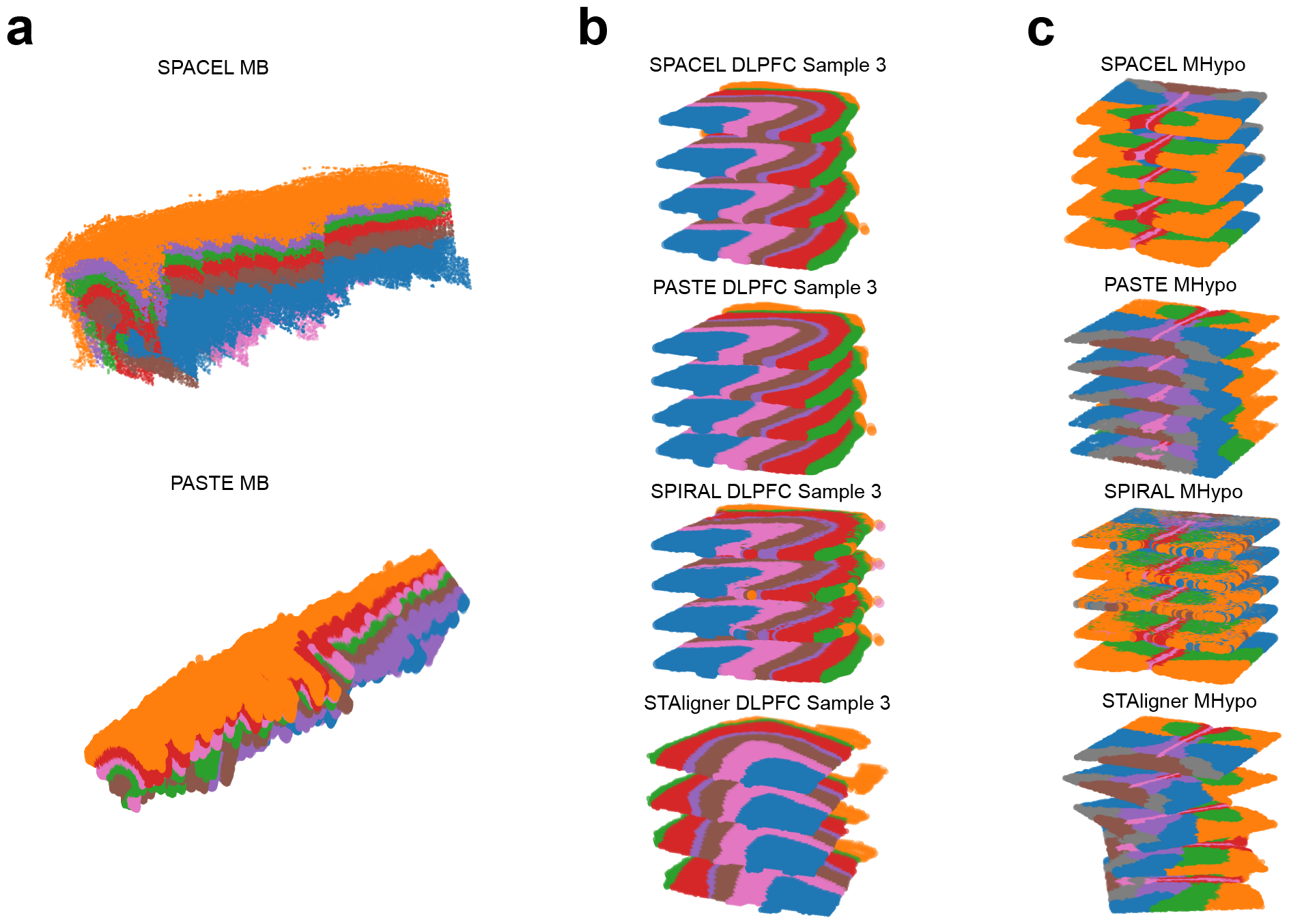
Reconstruction of 3D architecture of three different datasets. **a** 3D architecture reconstructed from 33 slices of MB data using SPACEL and PASTE. **b** 3D architecture reconstructed from four slices of DLPFC Sample 3 using SPACEL, PASTE, SPIRAL, and STAligner. **c** 3D architecture reconstructed from five slices of MHypo data using SPACEL, PASTE, SPIRAL, and STAligner.

In terms of the MB sample which contains 33 adjacent consecutive mouse brain tissue (Fig. 9c), only SPACEL and PASTE proved suitable for reconstructing the 3D architecture with this substantial number of slices. SPACEL successfully generated an effective 3D visualization. However, PASTE produced a discordant 3D architecture, particularly noticeable from the second half of the slices onward. Combining pairwise alignments from multiple adjacent slices into a stacked 3D alignment of tissue resulted in the propagation of errors. This explains the observation of two disjointed 3D architectures in the case of PASTE.

### Run time analysis for alignment and integration methods

Finally, we benchmarked the average runtime of each alignment and integration method on five selected datasets (Fig. 10). The two DLPFC datasets, 151673 and the anterior section of MB2SA&P were medium sized, with approximately 4k spots and 20-30k genes. Though each slice of the MHypo dataset has approximately 5k spots, each spot only contains 155 genes. The Embryo dataset is the largest in terms of the number of spots and genes. Lastly, the MB dataset has 33 slices in total. The plot of Fig. 10a, illustrates the average runtime when integrating two slices. Empty columns indicate scenarios where either the algorithm is not optimized for such use cases, or where memory consumption is excessively high, leading to the tool’s inability to complete execution. Overall, methods such as STAligner, PASTE, PASTE2, and PRE-CAST finished integration within 10 mins and exhibited reasonable scalability. Their time consumption was only marginally affected by increases in both the number of spots and genes. In contrast, scalability issues were more pronounced with methods like SPACEL, DeepST, and SPIRAL, where integration tasks might take hours or even days to complete. STAligner stands out as the sole tool capable of completing analysis on the Embryo dataset without encountering any memory issues thus far. In Fig. 10b, we further compared the runtime of each tool when integrating multiple (*>*2) slices. STAligner, PRECAST, and PASTE continued to exhibit promising scalability under these conditions. SPACEL and SPIRAL showed significantly slower performance, typically being 100x to 1000x slower than the aforementioned methods when integrating more than two slices. Notably, STAligner could finish on the MB dataset within 10 minutes.

**Figure 10.**
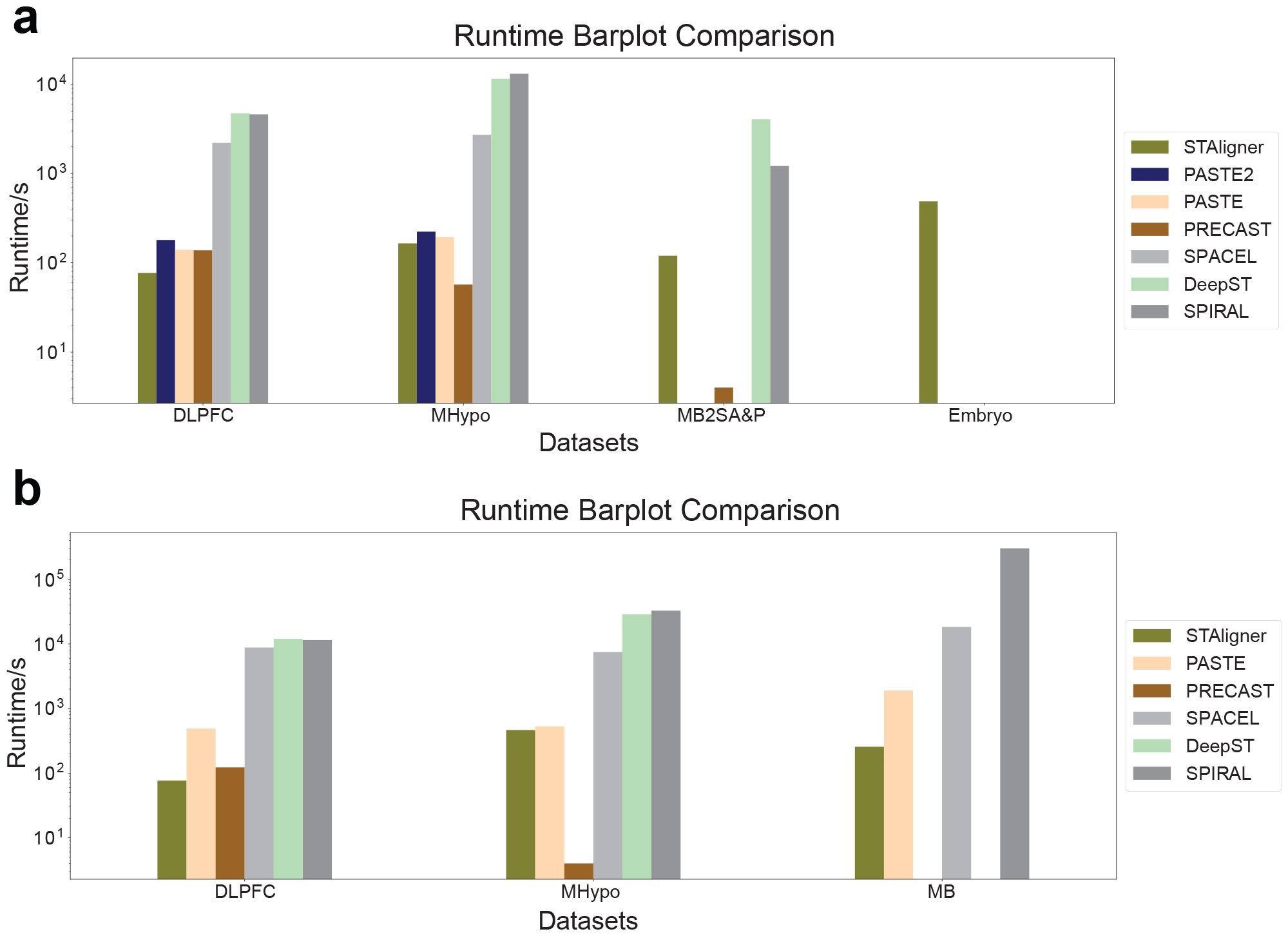
Comparison of runtime bar plots for different integration methods across five datasets. **a** Runtime for integrating two slices across four datasets. **b** Runtime for integrating multiple (*>*2) slices across three datasets. Empty columns for specific tools indicate scenarios where either the tool is not optimized for such use cases, or where the memory consumption is excessively high, resulting in the tool’s inability to complete execution.

## Discussion

In this study, we conducted comprehensive benchmark analyses covering different clustering, alignment, and integration tasks. We assessed 14 clustering methods, three alignment methods, and five integration methods across 68 slices of 10 publicly available ST datasets. We provided a user recommendation table 2 for users to choose an optimal tool to conduct corresponding analysis. We found that BASS, GraphST, ADEPT, and STAGATE outperformed the other 10 clustering methods. However, determining the definitive best-performing tool proved challenging, as each exhibited its highest performance across different datasets. Statistical methods consistently generated deterministic outputs with no variance in performance. In contrast, deep learning-based methods frequently yielded variance unless they employed identical seeds for each run. The overall performance trend for all methods decreased as the data complexity increased. All methods potentially suffer from algorithm overfitting, as indicated by their performance exceeding expectations on well-studied datasets but underperforming on less-studied ones. In terms of runtime, STAGATE, GraphST, and SpaGCN had the best scalability across datasets of different sizes.

**Table 2.** User recommendation table. This table ranks the top 3 most recommended tools for each analysis conducted.

| Analysis | top1 | top2 | top3 |
| --- | --- | --- | --- |
| Clustering accuracy | BASS | GraphST | ADEPT |
| Clustering robustness | BASS | ADEPT | GraphST |
| Integration (layer-wise alignment accuracy) | SPACEL | PASTE | STAligner |
| Integration (spot-to-spot mapping ratio) | PASTE | SPACEL | PRECAST |
| Integration on simulated data (spot-to-spot alignment accuracy) | PASTE2 | PASTE | SPACEL |
| Integration (joint embedding) | STAligner | DeepST | - |
| Joint clustering accuracy | BASS | STAligner | - |
| Integration (across conditions) | STAligner | BASS | - |
| 3D reconstruction | SPACEL | PASTE | - |
| Clustering Runtime | SpaGCN | GraphST | STAGATE |
| Integration Runtime | STAligner | PASTE | PRECAST |

While alignment and integration methods are capable of conducting multi-slice analysis, alignment methods such as PASTE, PASTE2, and SPACEL typically produce spot-to-spot alignment matrices or transformed spot coordinates based on alignment. In contrast, integration methods using deep learning backbones often generate joint spot embeddings for subsequent integration analyses. Therefore, it was not surprising to see that SPACEL and PASTE exhibited higher accuracy in layer-wise alignment compared to all integration tools as the primary objective of alignment methods was the direct alignment of spots across slices, rather than relying on joint spot embeddings for integration analysis. Relying on the joint spot embeddings to align spots across slices, STAligner achieved the highest layer-wise alignment accuracy among all integration methods, followed by DeepST, while PRE-CAST performed the least accurately. These results highlighted, to some extent, the inherent qualities of their learned joint spot embeddings. Our additional visualization plots for alignment-misalignment-unalignment analysis and spot-to-spot mapping ratios revealed that integration tools such as STAligner, SPIRAL, DeepST, and PRECAST produced joint spot embeddings capable of capturing global features for coarse layer-wise alignment and integration. Nevertheless, they might not suffice for capturing the local geometry necessary for spot-to-spot alignment. Our simulation experiments provided further validation for this observation. Notably, among all tools, PASTE2 and SPACEL achieved better spot-to-spot alignment accuracy when slices partially overlapped. The performance of all integration methods was highly sensitive to perturbation on the expression profiles.

Most integration methods were initially designed to learn joint spot embeddings across multiple slices. UMAP plots, projecting embeddings into two components, can to some extent reflect integration performance. Among these methods, STAligner stood out with better UMAP visualization, demonstrating integration with batch correction. However, its performance degraded for the MHypo datasets compared to the DLPFC datasets. SPIRAL, on the other hand, suffered from a significant batch effect due to uneven mixing of joint embeddings across slices, leading to notable separation issues across slices, consistent with its least favorable spot-to-spot mapping ratio. PRE-CAST tended to lose substantial geometry information by noticeably segregating spatial domains in the latent space, more so than the other tools. Although joint spot embeddings learned by multi-slice analysis have the potential to provide us a way to better detect spatial domains or cell types compared to single-slice analysis, it was difficult to conclude which tool had the overall best clustering performance in all pairs after joint training slices. In summary, there is still a need for a more robust integration tool. Integration methods could also align samples across different anatomic regions or development stages. We found STAligner outperformed other tools and had the scalability to process big datasets (over 50k spots).

As for the reconstruction of 3D architecture from multiple adjacent consecutive 2D slices, alignment tools such as PASTE and SPACEL outperformed integration tools like STAligner and SPIRAL. Specifically, in aligning a significant number of adjacent consecutive slices, SPACEL exhibited better performance than PASTE due to the potential for an erroneous alignment in PASTE to trigger a cascade of errors in subsequent slices. However, it is worth noting that the 3D reconstruction by SPACEL is not deterministic and exhibits variance. Nonetheless, we selected the most optimal 3D architecture for comparison purposes. Finally, in terms of runtime for alignment and integration, STAligner, PASTE, PASTE2, and PRECAST demonstrated good scalability for large datasets.

## Methods

### Quantitative analysis for clustering

#### 1. Benchmark metrics

- Adjusted Rand Index (ARI): ARI is a measure of the similarity between two data clusterings. It is a correction of the Rand Index, which evaluates the concordance between pairs of data points, determining whether they are grouped together or separated in two different clusterings. The ARI value is calculated using Equations 1 and 2. *a* is the number of pairs of elements that are in the same cluster in both the ground true and predicted clusterings, *b* is the number of pairs of elements that are in different clusters in both the ground true and predicted clusterings, *c* is the number of pairs of elements that are in the same cluster in the true clustering but in different clusters in the predicted clustering, and *d* is the number of pairs of elements that are in different clusters in the true clustering but in the same cluster in the predicted clustering. *Expected RI* is the expected value of the Rand Index under the assumption of independence between the true and predicted clusterings. *Max RI* is the maximum possible Rand Index.
- Spatial Coherence Score (SCS): A spatial coherence score of the cluster labels is computed based on O’Neill’s spatial entropy. A high spatial coherence score indicates that the cluster labels of adjacent spots are frequently identical, while a low spatial coherence score suggests that cluster labels of adjacent spots are drawn from the uniform distribution. This score serves as an indicator of data quality. Specifically, let *G* = (*V, E*) be a graph where *V* is the set of spots, and edges (*i, j*) ∈ *E* connect every pair (*i, j*) of adjacent spots. Let *K* = *{*1, 2, …, *k}* be a set of *k* cluster labels, and let *L* = [*l*(*i*)] be a set of labelings of spots, where *l*(*i*) ∈ *K* is the cluster label of spot *i* The spatial entropy *H*(*G, L*) is defined in Equation 3, where 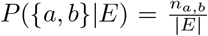, and *n*_*a,b*_ is the number of edges (*i, j*) ∈ *E* such that *l*(*i*) = *a* and *l*(*j*) = *b*. The spatial coherence score is defined as a normalized form of spatial entropy, using the absolute value of the Z score of spatial entropy over random permutations of the labels of spots in a slice [31].
- Runtime: We collected the average runtimes from 20 iterations for each clustering method across all benchmarking datasets to assess their scalability.

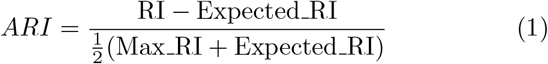

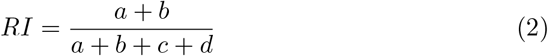

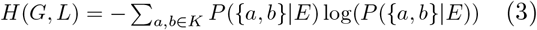

#### 2. Domain identification performance across 31 ST slices

Given that spatial domain or cell type identification is the primary objective of clustering methods, we aim to conduct a thorough performance comparison using ARI when manual annotation serving as ground truth is available. One notable challenge for deep learning methods is the variance introduced by random seed initialization. To address this, we computed the average ARI from 20 runs on each dataset and displayed these results using boxplots and a heatmap plot to enhance comparison and visualization. Additionally, since there are 31 ST slices across 7 different datasets, it is challenging to rank the overall performance solely based on the average ARI heatmap plot. Therefore, we also provided another heatmap for the overall ranking. This ranking heatmap was generated by normalizing all results within the same slice by dividing them by the maximum ARI value (representing the best performance) among all methods, thereby standardizing all ARI values to 1. With 31 data slices in total, for each method, the best ranking for the sum result is 31, while the best ranking for the average result is 1. To ensure fairness, the rank scores were averaged exclusively over feasible ST data, excluding instances with NaN values.

#### 3. Overall robustness across seven ST datasets

To assess the robustness of methods on each dataset, the clustering results across different ST slices within the same dataset were averaged. A robust method is expected to demonstrate the highest overall ARI value across all datasets, even if it may encounter challenges in predicting a few individual slices.

#### 4. Data complexity effect on method performance

Data complexity is recognized to have an impact on method performance. Although different methods are often fine-tuned on different datasets to demonstrate superiority in specific contexts, our objective is to identify a general trend wherein methods exhibit diminished performance as data complexity increases. In this context, the Spatial Coherence Score (SCS) is introduced as a metric for quantifying data complexity. The underlying assumption is that data with more coherent regions, indicated by a higher SCS, are easier for domain identification.

### Qualitative analysis for clustering

#### 1. Clustering evaluation by visualization

For MHPC data without region-based annotation, the evaluation is constrained to comparing the clustering results with the cell type annotation through visualization, supplemented by reference to the Mouse Allen Brain atlas.

### Quantitative analysis for batch correction and integration

#### 1. Benchmark metrics

- Adjusted Rand Index: As illustrated in the clustering metrics section.
- Layer-wise alignment accuracy: This metric relies on an important hypothesis that aligned spots from adjacent consecutive slices within a dataset are more likely to pertain to the same spatial domain or cell type. Joint spot embeddings learned from each method are utilized to align (anchor) spots from the first slice to (aligned) spots on the second slice for each slice pair. This alignment accuracy is defined as the ratio of the number of anchor spots to the total number of spots within the first slice when anchor spots and aligned spots belong to the same spatial domain or cell type. Euclidean distance is employed to define the closeness of spots to be aligned. A good integration tool is expected to demonstrate high accuracy for anchor and aligned spots belonging to the same spatial domain or cell type.

For DLPFC data which has a unique layered structure, this metric is also meticulously designed to demonstrate whether anchor and aligned spots belong to the same layer (layer shifting = 0) or they belong to different layers (layer shifting = 1 to 6).

- Spot-to-spot matching ratio: This metric further evaluates whether joint embeddings’ quality captures the data geometry. The ratio is defined as the ratio of the total number of anchor spots from the first slice to the number of aligned spots from the second slice. For two adjacent consecutive slices, a nearly 1:1 ratio is expected for an optimal tool.
- Spot-to-spot alignment accuracy: This metric is used to evaluate joint embeddings for simulated datasets since the ground truth for spot-to-spot alignment relationship is available. This spot-wise alignment accuracy is defined as the percentage of anchor spots from the first slice that match correctly to aligned spots on the second slice.

#### 2. Comparison of clustering performance before and after integration

A good practice that connects integration and clustering tasks is multi-slice joint clustering. To determine if incorporating information from adjacent consecutive slices enhances domain or cell type identification, we used batch-corrected joint embeddings to evaluate clustering results on each single slice based on ARI values. We plotted ARI for clustering results before and after the integration. However, some integration methods do not support single-slice clustering. We thus only plotted ARI after the integration of these methods.

#### 3. Simulated data for integration

Given the scarcity of benchmark datasets available for integration tasks to evaluate spot-to-spot alignment accuracy, we modified the simulation method proposed in PASTE [31] and generated 11 simulated 10x Visium datasets for this evaluation. We first used one DLPFC slice (151673) as the reference and simulated additional slices with different overlapping ratios (20%, 40%, 60%, 80%, and 100%) in comparison to the reference slice. In this simulation scenario, the pseudo-count perturbation was fixed at 1.0 for all simulated slices. Next, we simulated additional slices with different pseudocounts (0 - 3.0 with a step size of 0.5) to represent perturbation on gene expression while keeping the overlapping ratio fixed at 80%. Specifically, by taking the DLPFC 151673 slice as the reference, we altered the spatial coordinates in the new slice by rotating this reference slice, perturbed the gene expression by adding pseudocounts, and adjusted the number of spots by removing some spots that did not align with the grid coordinates following the rotation. To keep fidelity with the real 10x Visium data, the spots within the tissue in our simulation are arranged in a hexagonal grid rather than in a rectangular grid pattern. Additionally, we utilized the minimal distance between adjacent spots on the DLPFC 151673 slice as the distance between any two adjacent simulated spots on the grid, rather than arbitrarily setting it to 1.

More detailed procedures to generate simulated datasets are described as follows.

- Create a hexagonal grid *G*. Let *g*_.*i*_ and *z*_.*k*_ denote the 2D coordinates of spot *i* on grid *G* and spot *k* on the reference slice DLPFC 151673, respectively. *d*_*ij*_ = ||*g*_.*i*_ *− g*_.*j*_|| = min_*kl*_ ||*z*_.*k*_ *− z*_.*l*_|| for any two adjacent simulated spots *i* and *j* on grid that *i, j* ∈ *G*.
- Let *R* be a rotation matrix with an angle *θ*. After spot *k* is rotated with an angle *θ*, the rotated coordinates of spot *k, r*_.*k*_ = *Rz*_.*k*_, is used to mapped the spot *k* to the closest grid spot *î* by *î* = arg min_*i*_ ||*g*_.*i*_ *− r*_.*k*_||. Then, the simulated coordinates of tissue spot *k*, 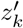 is given by 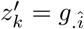 Spot *k* is dropped if the grid spot *g*_·*î*_ was already used by a previous tissue spot.
- Let *X* = [*x*_*ij*_] ∈ ℕ^*m×n*^ represent the *m* genes by *n* spots expression profile matrix of DLPFC slice 151673, where *x*_*ij*_ is the read count of gene *i* in tissue spot *j*. We can calculate the mean of the total transcript count of the tissue spots, 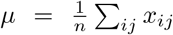 and the variance of the total read count,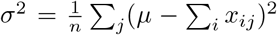. Total read counts of spot *j, k*_*j*_, are generated according to *k*_*j*_*∼*NegativeBinomial(*r, p*). Here 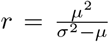 and 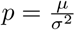 such that *E*(*k*_*j*_) = *µ* and var(*k*_*j*_) = *σ*^2^.
- Generate simulated gene *i* read count for spot *j* according to 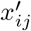 *∼*Multinomial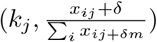, where *δ* ∈ {0, 0.5, …, 3} is a pseudocount.

### Qualitative analysis for batch correction and integration

#### 1. Visualization of aligned, misaligned, and unaligned spots from pairwise alignment

To assess the joint spot embeddings by integration tools and the alignment matrices by alignment tools, we quantified the alignment accuracy based on aligned, misaligned, and unaligned spots across two consecutive slices. For integration tools such as STAligner, PRE-CAST, DeepST, and SPIRAL, we aligned the spot (referred to as the “anchor” spot) on the first slice with the spot (referred to as the “aligned” spot) on the second slice based on their joint latent embeddings using Euclidean distance. If the aligned spot belonged to the same spatial domain or cell type as the anchor spot according to ground truth labels, we classified both spots as “aligned” spots (denoted as “orange” color in Fig. 4c-d). If the aligned spot did not belong to the same spatial domain or cell type as the anchor spot, we classified both spots as “misaligned” spots (denoted as “blue” color in Fig. 4c-d). In the last scenario, if spots on the second slice were not used to match any spot on the first slice, these spots on the second slice were classified as “unaligned” spots (denoted as “green” color in Fig. 4c-d). For alignment tools like PASTE, PASTE2, and SPACEL, we directly used their alignment matrices to perform this analysis.

#### 2. Reconstruction of three-dimensional (3D) architecture of the tissue

Among all integration methods, there are only three tools, PASTE, PASTE2, and SPACEL that have an output for a transformed coordinate system for all slices. Consequently, they can combine pairwise alignments from multiple adjacent consecutive slices into a stacked 3D alignment of a tissue. These three tools were benchmarked in three datasets by comparing their 3D architecture of the tissue.

#### 3. Visualization of UMAP plot for joint embeddings

Most integration methods primarily concentrate on embedding the spots within a high-dimensional latent space, which often proves challenging to interpret intuitively. To enhance comprehension of the distribution in the latent space, we performed dimension reduction for spot embeddings to two dimensions using UMAP. A quality UMAP plot of latent embeddings should exhibit structures resembling those of the real data while also demonstrating spatial domain or cell types in a separable manner.

#### 4. Visualization of clustering results after integration

For the MB2SA&P dataset, we compared the identified domains after integration with the Allen Brain atlas through visualization. Furthermore, we examined the consistency of regions across the fissure between the anterior and posterior sections. Higher similarity to the atlas, along with the region coherence, serve as indicators of superior integration performance.

For the mouse Embryo data, we compared the clustering result after integrating two slices for developmental stages E11.5 and E12.5 with the manual annotation defined by different organs and tissues.

### Computation platform

We conducted all benchmarking experiments on our computer server equipped with one Intel Xeon W-2195 CPUs, running at 2.3 GHz, featuring a total of 25 MB L3 cache, and comprising 36 CPU cores. The cluster also boasted 256 GB of DDR4 memory operating at 2,666MHz.

For the GPU configurations, we utilized the same computer with four Quadro RTX 8000 cards, each having 48 GB of memory and a total of 4608 CUDA cores.

## Availability of data and materials

All code, tutorials, and related data files are available at https://github.com/maiziezhoulab/BenchmarkST and https://benchmarkst-reproducibility.readthedo4c.s.io/en/latest/. All data and the corresponding annotation can be downloaded from https://benchmarkst-reproducibility.readthedocs.io/en/latest/Data%20availability.html and are described in Table 1 with their sources. Dataset 1 consists of 12 human DLPFC sections with manual annotation [39]. Dataset 2 includes a single slice of human breast cancer, which is open-sourced from 10x genomics [22]. Dataset 3 includes two slices of anterior and posterior mouse brain [18]. Dataset 4 contains HER2-positive tumors from eight individuals [40]. Dataset 5, including anatomical regions of the mouse hippocampus, is acquired through the Broad Institute [41]. Dataset 6 is the Embryo dataset sequenced by Stereo-seq from the MOSTA project [42]. Dataset 7 contains one slice from the mouse visual cortex [9]. Dataset 8 contains three slices of the mouse prefrontal cortex [9]. Dataset 9 includes five slices from the mouse hypothalamus [18]. Dataset 10 contains 33 consecutive mouse cerebral cortex tissue slices with similar shapes [33].

The simulation data is deposited in Zenodo https://zenodo.org/records/10800745.

## Funding

This work was supported by the NIGMS Maximizing Investigators’ Research Award (MIRA) R35 GM146960 to X.M.Z., and Guangdong Basic and Applied Basic Research Foundation (2023A1515030154) to W.S..

## Competing interests

The authors declare that they have no competing interests.

## Author’s contributions

X.M.Z. conceived and led this work. Y.H. and X.M.Z. designed the framework. Y.H., Y.L., M.X. M.R. W.S., C.L., H.Q., and J.B. performed all benchmark analyses. Y.H. and X.M.Z. wrote the manuscript with input from all authors.

## Notes

### Competing Interest Statement

The authors have declared no competing interest.

