## Appendix for "Benchmarking clustering, alignment, and integration methods for spatial transcriptomics"

Hu et al.

### 1 Supplementary Figures

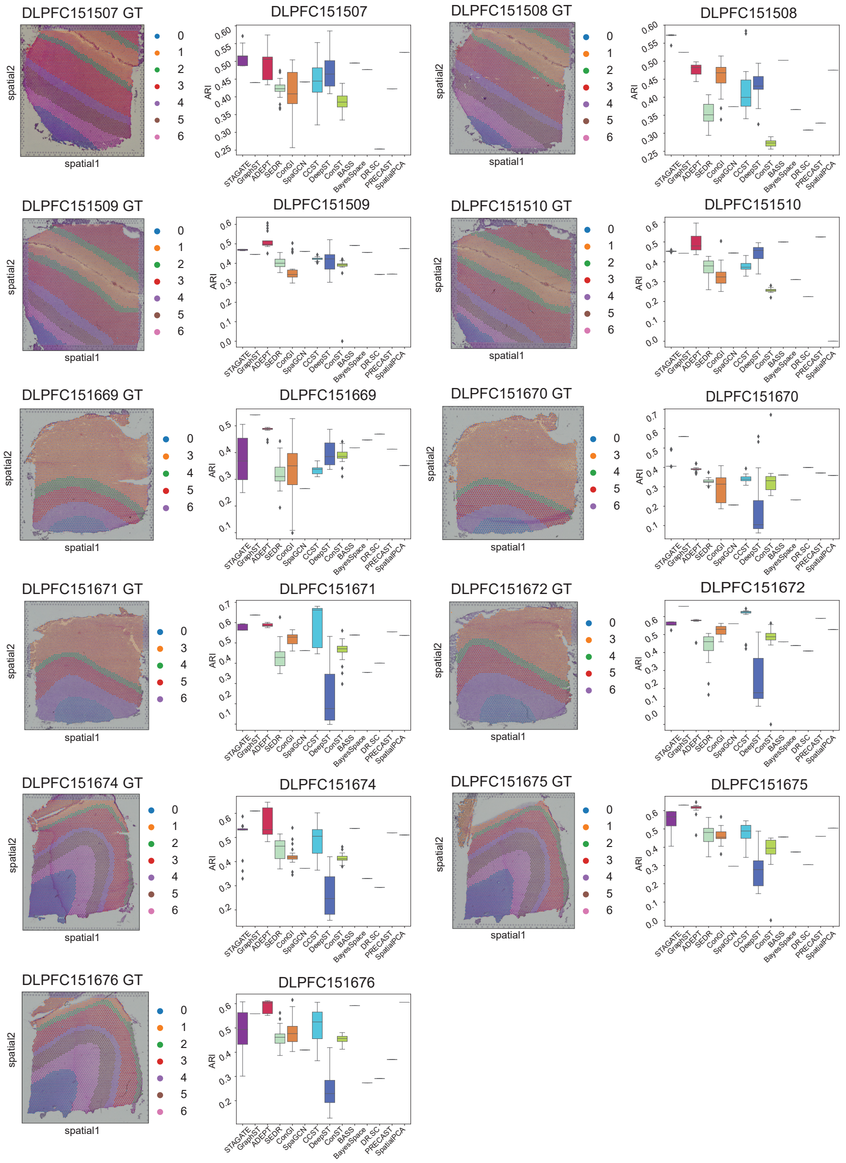

Figure S1: **The ground truth visualization plots and ARI box plots.** The ground truth visualization plots and box plots depicting ARI values of all tools on 11 DLPFC slices.

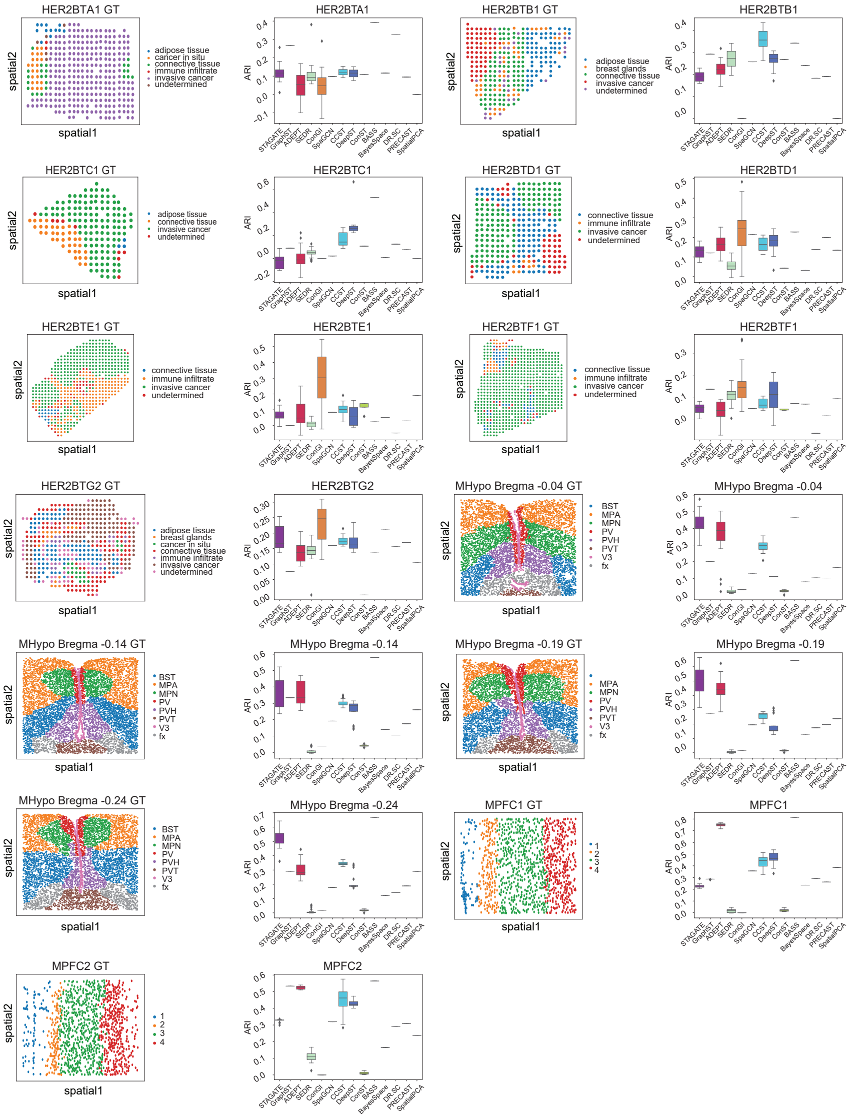

Figure S2: **The ground truth visualization plots and ARI box plots.** The ground truth visualization plots and box plots depicting ARI values of all tools on seven HER2BT slices, four MHypo slices, and two MPFC slices.

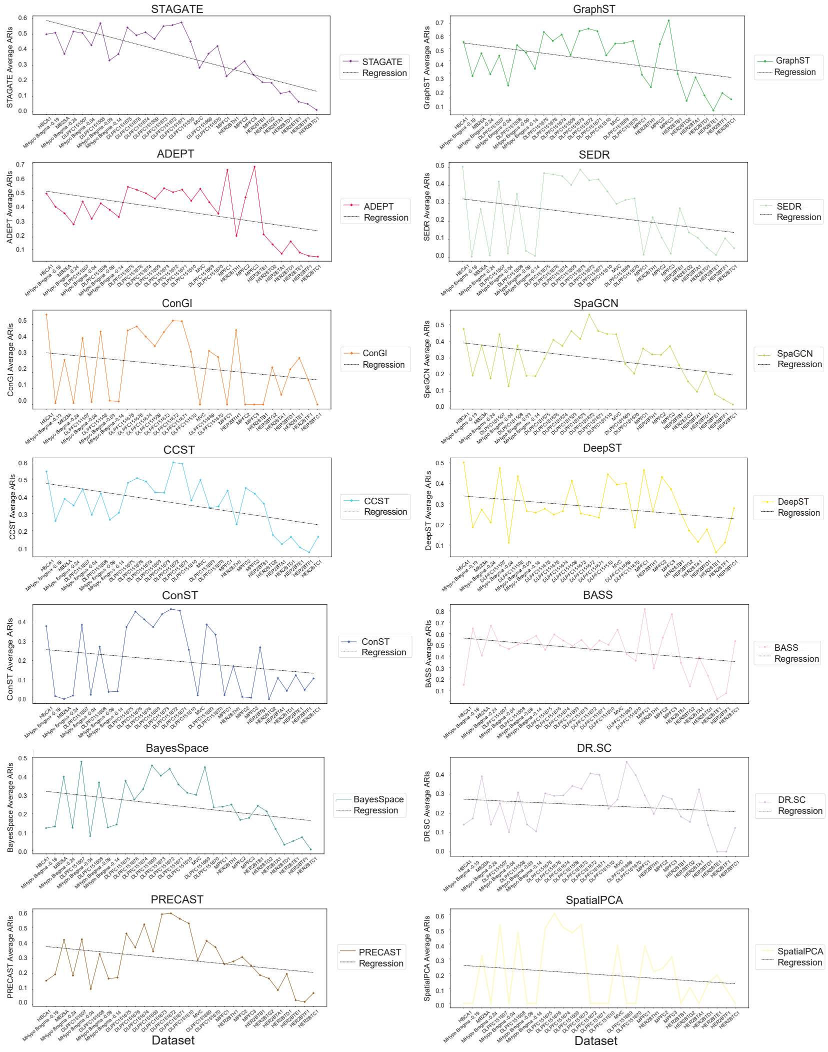

Figure S3: **ARI values vs. data complexity for each clustering tool**. Each line plot with a regression line shows the decline in average ARI for each method as data complexity increases across 31 ST slices.

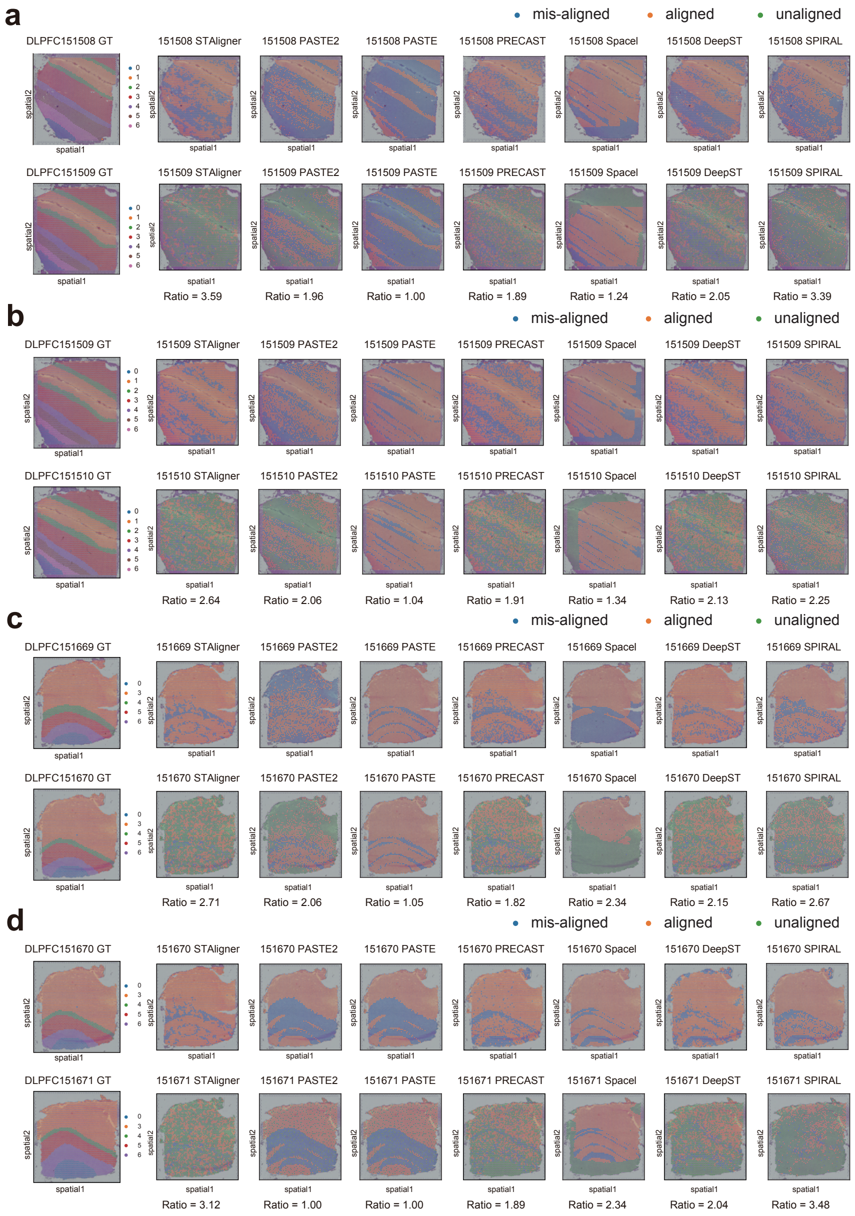

Figure S4: **Visualization plots for alignment-misalignment-unalignment for the DLPFC dataset.** These visualization plots show aligned spots, misaligned spots, and unaligned spots when aligning the anchor spot from the first (top) slice to the aligned spots on the second (bottom) slice on DLPFC 151508-151509, 151509-151510, 151669-151670, and 151670-151671 pairs. Values below each plot represent the spot-to-spot matching ratio.

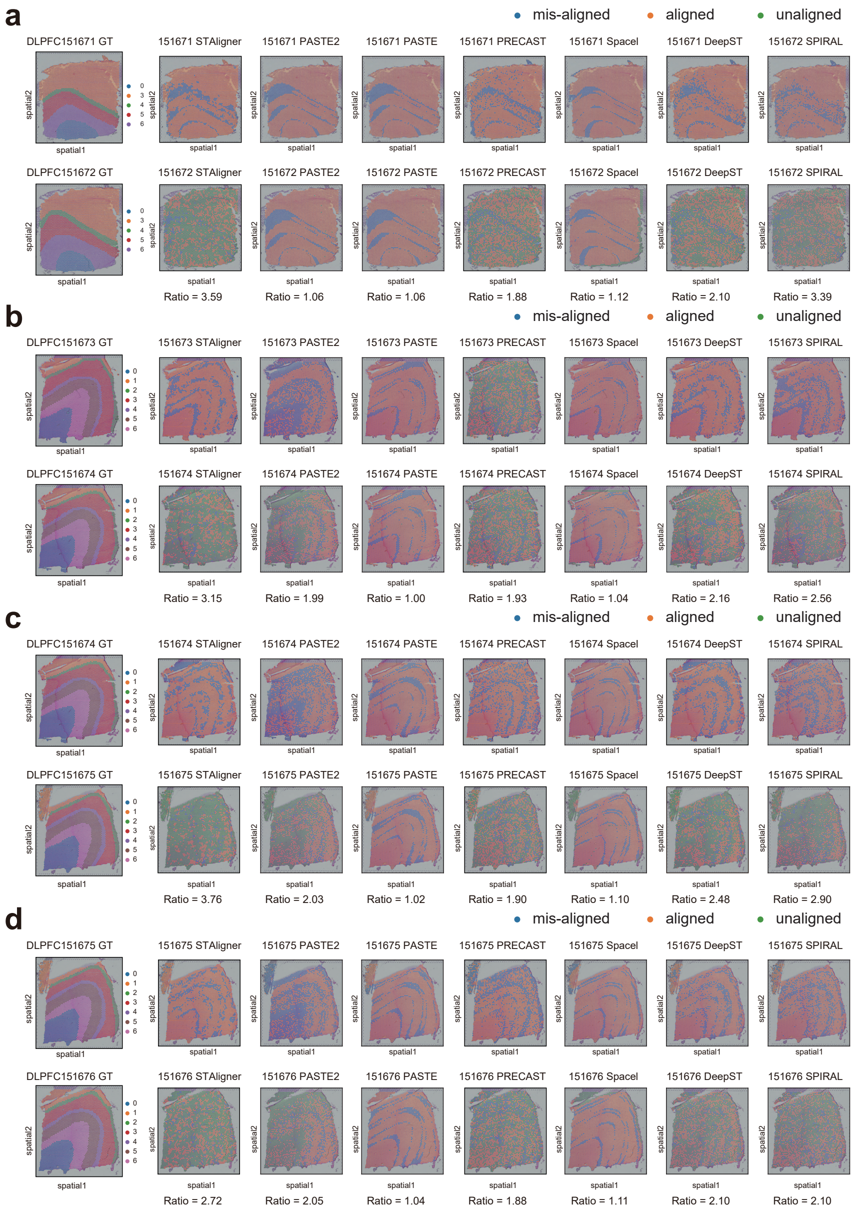

Figure S5: **Visualization plots for alignment-misalignment-unalignment for the DLPFC dataset.** These visualization plots show aligned spots, misaligned spots, and unaligned spots when aligning the anchor spot from the first (top) slice to the aligned spots on the second (bottom) slice on DLPFC 151671-151672, 151673-151674, 151674-151675, and 151675-151675 pairs. Values below each plot represent the spot-to-spot matching ratio.

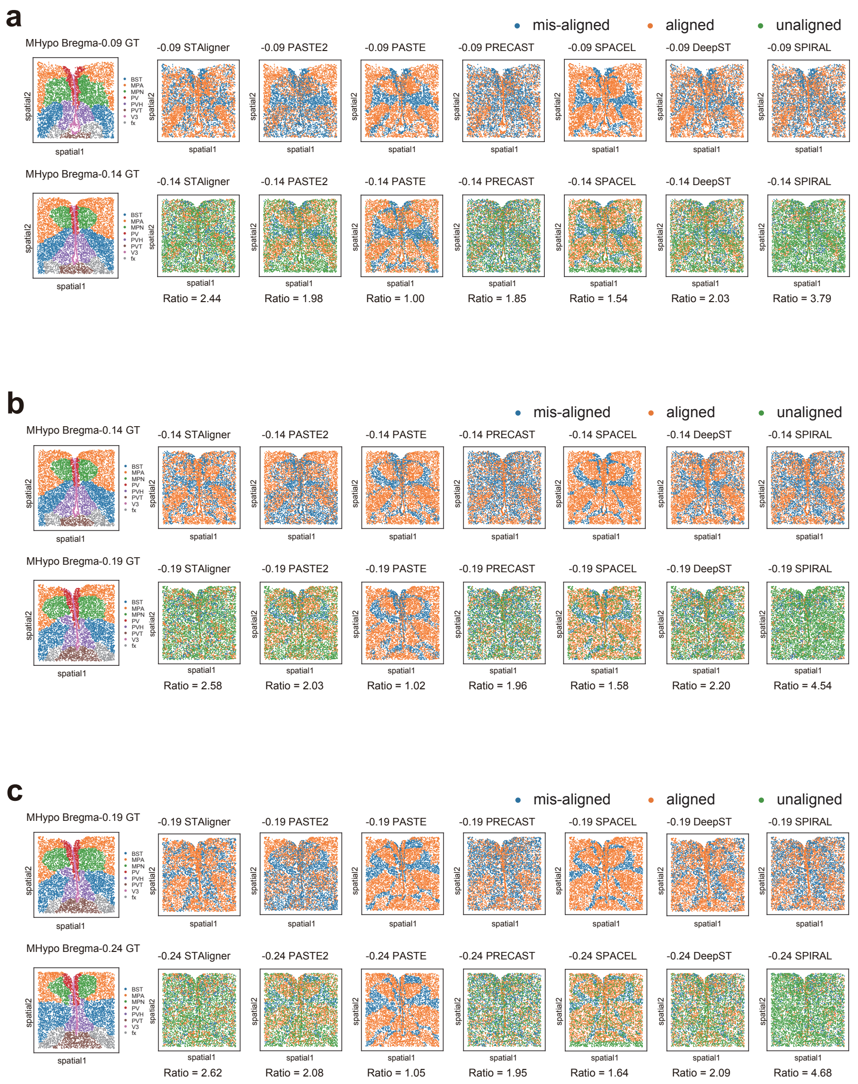

Figure S6: **Visualization plots for alignment-misalignment-unalignment for the MHypo dataset.** These visualization plots show aligned spots, misaligned spots, and unaligned spots when aligning the anchor spot from the first (top) slice to the aligned spots on the second (bottom) slice on MHypo Bregma -0.09 - -0.14, -0.14 - -0.19, and -0.19 - -0.24 pairs. Values below each plot represent the spot-to-spot matching ratio.

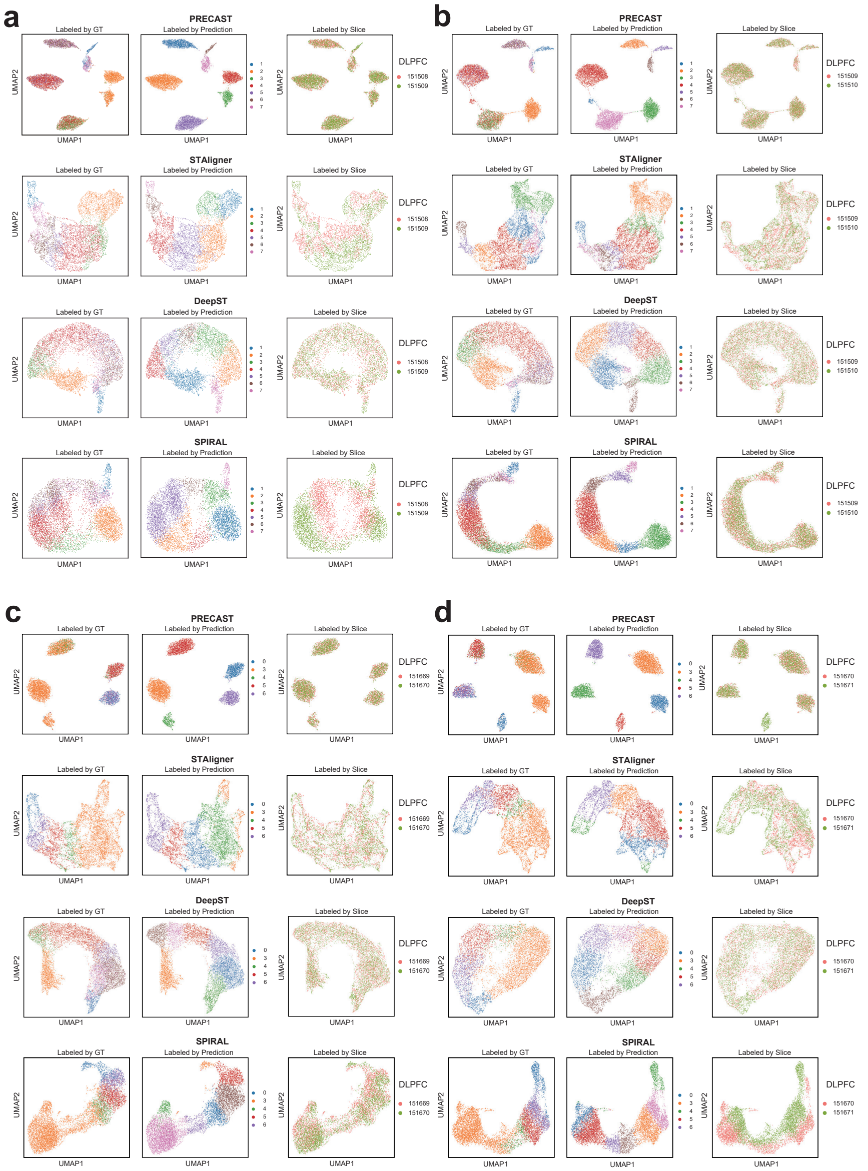

Figure S7: **UMAP plots of low dimensional joint embedding distribution for batch correction for the DLPFC dataset. a-d** These UMAP plots depict the 2D distribution of latent joint embeddings after integration with batch correction by different integration methods on the DLPFC 151508-151509 pair (a), the DLPFC 151509-151510 pair, **(c)** the DLPFC 151669-151670 pair, and **(d)** the DLPFC 151670-151671 pair. Each UMAP contains colored spots labeled by three different setups: ground truth (GT), method prediction, and slice index.

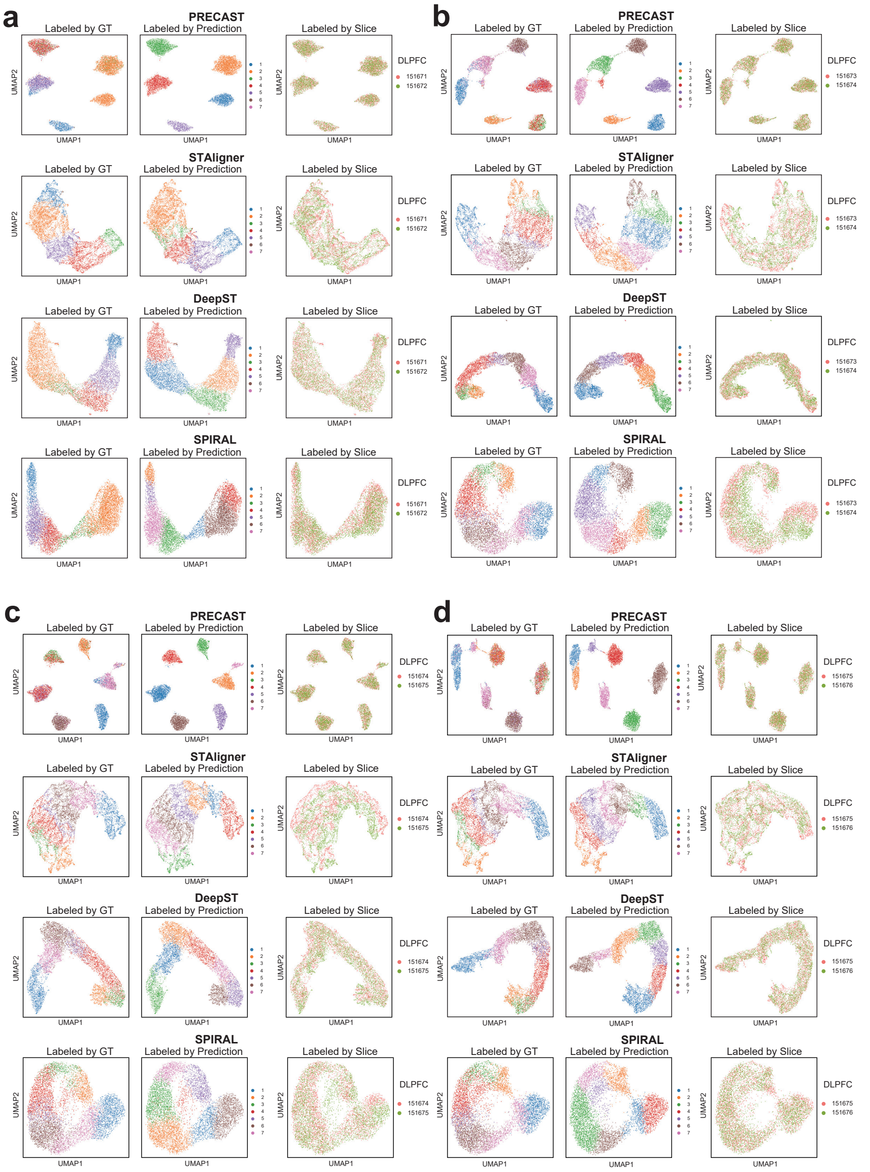

Figure S8: **UMAP plots of low dimensional joint embedding distribution for batch correction for the DLPFC dataset. a-d** These UMAP plots depict the 2D distribution of latent joint embeddings after integration with batch correction by different integration methods on the DLPFC 151671-151672 pair (a), the DLPFC 151673-151674 pair (b), the DLPFC 151674-151675 (c), and the DLPFC 151675-151676 pair (d). Each UMAP contains colored spots labeled by three different setups: ground truth (GT), method prediction, and slice index.

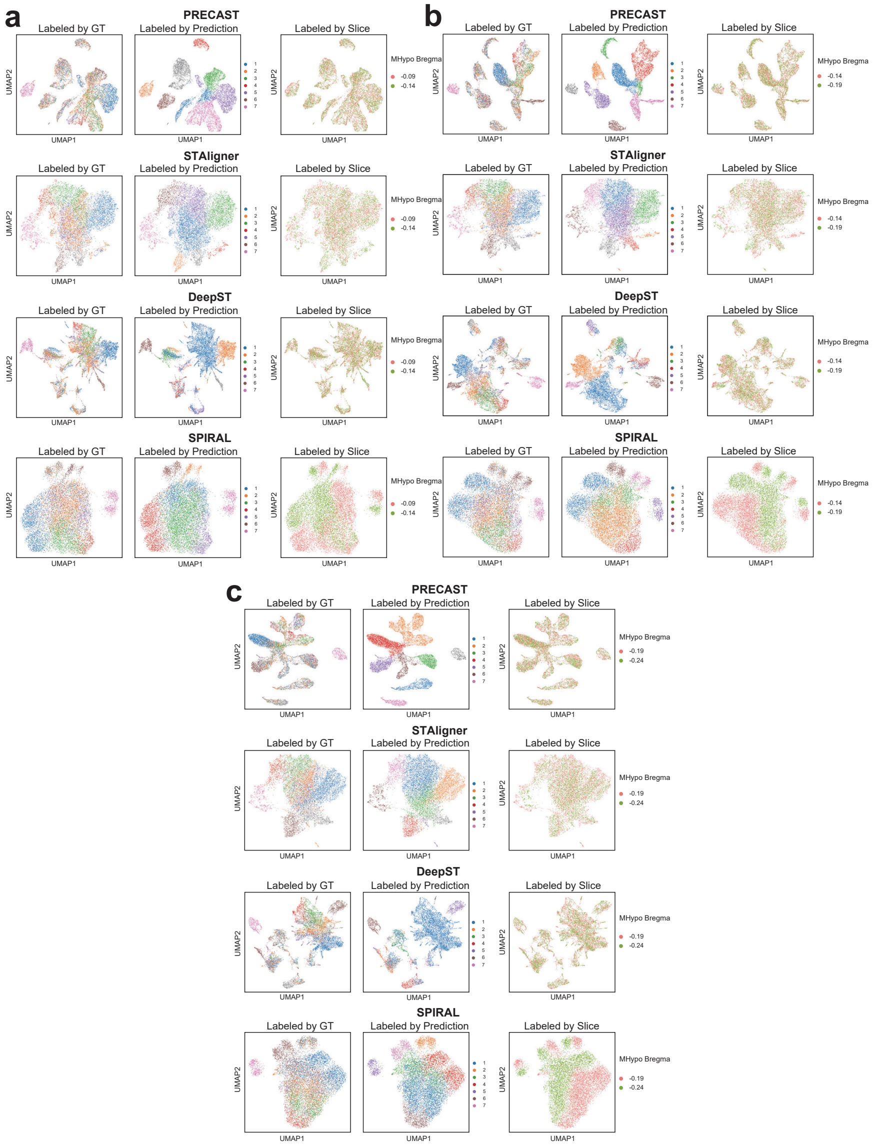

Figure S9: **UMAP plots of low dimensional joint embedding distribution for batch correction for the DLPFC dataset. a-d** These UMAP plots depict the 2D distribution of latent joint embeddings after integration with batch correction by different integration methods on the MHypo Bregma -0.09 - -0.14 pair (a), the MHypo Bregma -0.14 - -0.19 pair (b), and the MHypo Bregma -0.19 - -0.24 pair (c). Each UMAP contains colored spots labeled by three different setups: ground truth (GT), method prediction, and slice index.

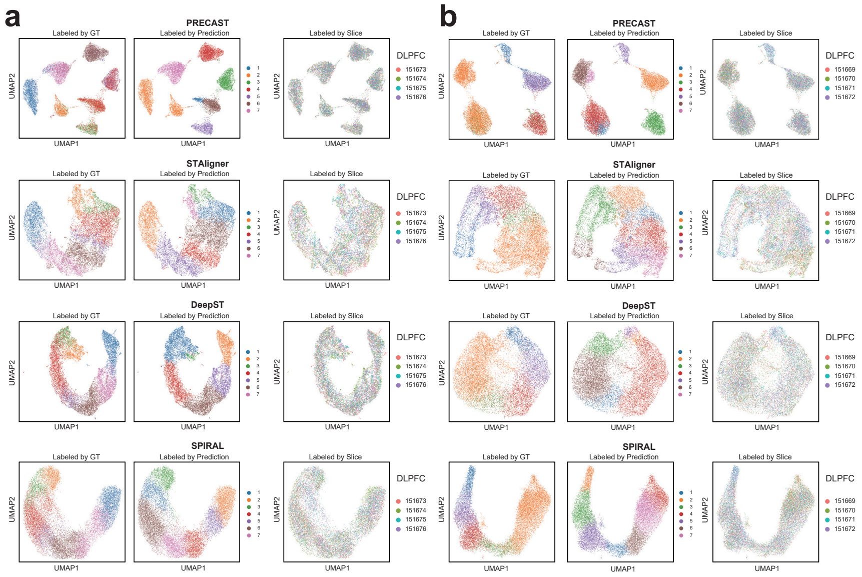

Figure S10: **UMAP plots of low dimensional joint embedding distribution for batch correction. a-b** These UMAP plots depict the 2D distribution of latent joint embeddings after integration with batch correction by different methods on the DLPFC 151673-151676 four slices (a), and the DLPFC 151669-151672 four slices (b). Each UMAP contains colored spots labeled by three different setups: ground truth (GT), method prediction, and slice index.
